# Degradation of the TEAD•YAP/TAZ Transcription Factor Complex by Heterobifunctional Small Molecules that Bind to the TEAD Allosteric Lipid Pocket

**DOI:** 10.64898/2026.01.09.698724

**Authors:** I-Ju Yeh, Mona K. Ghozayel, Khuchtumur Bum-Erdene, Samy O. Meroueh

**Affiliations:** Department of Biochemistry, University of Illinois Urbana-Champaign, Urbana, IL, 61801, USA; Cancer Center at Illinois, University of Illinois Urbana-Champaign, Urbana, IL, 61801, USA; Department of Veterans Affairs, Richard L. Roudebush VA Medical Center, Indianapolis, Indiana, 46202; Department of Biochemistry and Molecular Biology, Indiana University School of Medicine, Indianapolis, IN, 46202, USA

## Abstract

Central kinases of the Hippo tumor suppressor pathway phosphorylate the transcriptional coactivators YAP and TAZ to sequester them in the cytoplasm. In cancer, Hippo pathway kinases have reduced activity, leading to translocation of YAP and TAZ into the nucleus where they engage TEADs and other transcription factors. Here, we explore whether heterobifunctional small molecules that bind to the TEAD allosteric lipid-binding pocket can degrade the TEAD•YAP/TAZ complex. We design and synthesize heterobifunctional molecules that consist of flufenamic acid analogs that bind to the allosteric TEAD lipid pocket, a long and flexible linker, and thalidomide to engage E3 ubiquitin ligase component cereblon. The bifunctional compounds promote ternary complex formation in biochemical assays and mammalian cells but exhibited modest degradation of TEAD, YAP, and TAZ in cancer cells. Methyl ester analogs of these compounds led to substantial proteasomal degradation of the TEAD•YAP/TAZ complex in cancer cells. This work provides a strategy for depletion of nuclear YAP and TAZ and for exploration of their TEAD-dependent and TEAD-independent activities *in vivo*.

## INTRODUCTION

Transcriptional regulator yes-associated protein (YAP) or its paralog transcriptional coactivator with PDZ-binding motif (TAZ) binds to transcriptional enhancer associated domain (TEAD) transcription factors to form a protein-protein complex that regulates Hippo pathway gene expression (1, 2). The core of the Hippo pathway includes two tumor suppressor kinases, large tumor suppressor kinases 1 (LATS1) and 2 (LATS2), and mammalian STE20-like protein kinases 1 (MST1) and 2 (MST2), which phosphorylate YAP and TAZ. Phosphorylated YAP and TAZ are either bound to 14-3-3 proteins in the cytoplasm or degraded in the proteasome (3). When Hippo kinases are inactivated, unphosphorylated YAP and TAZ are translocated to the nucleus, where they bind to TEADs and other transcription factors such as AP-1 and RUNX. These transcriptional complexes regulate the expression of *CTGF*, *Cyr61*, *Myc*, *WNT5A/B*, *DKK1* and others (4–14).

Over the past two decades, a large number of studies on the Hippo pathway have revealed that YAP and TAZ play important roles in tumor formation, growth, and metastasis in nearly every tissue type (15–23), as described in two recent reviews (24, 25). The roles of YAP and TAZ in cancer are attributed to a wide range of cellular cues and receptors that control Hippo signaling, including G-protein coupled receptors (GPCRs), alterations in tight and adherens junctions (26), energy stress (27), osmotic stress, heat shock, fluctuations in glycogen levels (28), and mechanical forces (29). In hematologic cancers, YAP and TAZ are downregulated, but their roles are complex and context dependent as they have been shown to be either oncogenic or tumor suppressive.

There is substantial interest in the development of small molecules that directly inhibit YAP and TAZ. This is highly challenging as YAP and TAZ lack well-defined binding pockets. The most common strategy for inhibiting YAP and TAZ is to target TEAD transcription factors (30–34). A large number of small-molecule covalent and non-covalent TEAD antagonists have been developed over the past five years. Covalent inhibitors form a covalent bond with a conserved cysteine residue in the binding pocket. Most of these compounds do not disrupt the TEAD•YAP/TAZ interaction, because their binding site is located outside the protein-protein interface. However, a few compounds were shown to inhibit this interaction (35). Despite the high affinity and large interface between YAP/TAZ and TEAD, small molecules that bind at the interface have been recently reported (36). Also, proteolysis-targeting chimeras (PROTACs) have been developed to degrade TEAD transcription factors (37, 38).

Here, we develop proteolysis-targeting chimeras (PROTACs) that bind to the TEAD allosteric pocket and achieve proteasomal degradation of the TEAD•YAP/TAZ complex. The bifunctional compounds consist of a TEAD flufenamic acid analog ligand, a linker, and the cereblon ligand thalidomide. Binding of the bifunctional molecule to the TEAD allosteric pocket resulted in the degradation of TEAD, YAP, and TAZ. We designed and synthesized several bifunctional molecules that led to the proteasomal degradation of all three proteins in cancer cells.

## RESULTS

### Design and Synthesis of Heterobifunctional Compounds

An X-ray crystal structure of the drug flufenamic acid bound to the TEAD palmitate binding pocket (PDB ID: 5DQ8) provided a starting point for the design of heterobifunctional small molecules as potential PROTAC degraders [**Fig. 1A**]. The structure revealed that the *ortho* position relative to the carboxylic acid moiety of the drug was located near the opening of the palmitate binding pocket. We chose this position to attach a linker. We also used the above X-ray structure to design derivatives of flufenamic acid to enhance its binding affinity to the TEAD lipid pocket. Superimposition of the TEAD-flufenamic acid X-ray structure with a TEAD-palmitate X-ray structure revealed that palmitate binds deeper into the pocket and extends beyond the trifluoromethylphenyl moiety of flufenamic acid (**Fig. 1A**). Based on this, we designed five analogs of flufenamic acid that possess different substituents at R_2_ to mimic the deep-pocket aliphatic chain of palmitate (**Fig. 1B**). Finally, a 15-atom linker was designed to tether the flufenamic analogs to the cereblon ligand, thalidomide, thereby generating several heterobifunctional molecules: Carboxylic acids **1** (TED-651), **2** (TED-677), **3** (TED-672), **4** (TED-675), and **5** (TED-673); and methyl esters **6** (TED-650)**, 7** (TED-688)**, 8** (TED-690)**, 9** (TED-689), and **10** (TED-734) (**Fig. 1B**). The esters were prepared using a prodrug strategy to enhance cell permeability of the carboxylic acid compounds, since the bifunctional molecules must cross the plasma membrane and localize to the nucleus to reach the intended protein targets.

**Figure 1.**
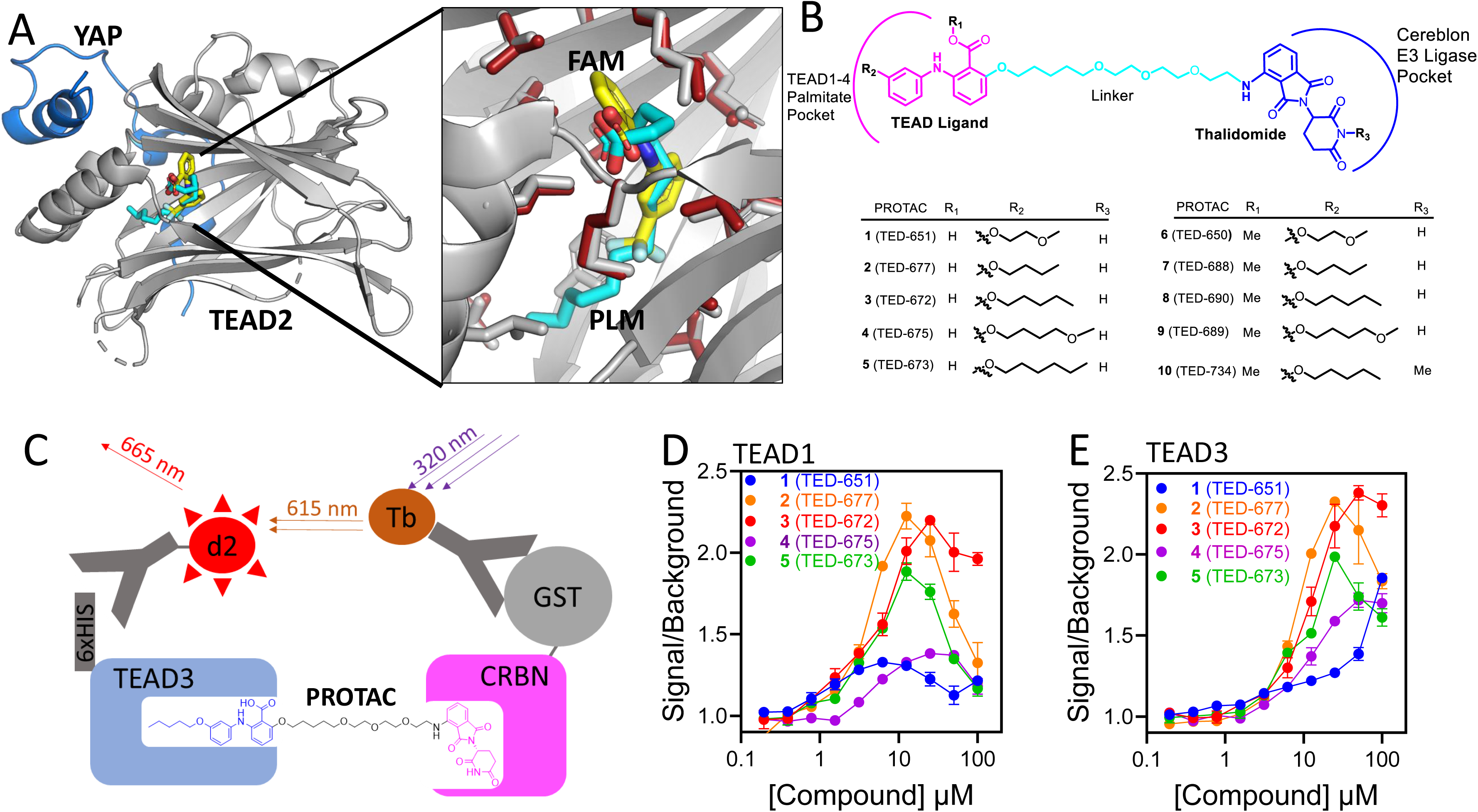
PROTAC Design and Biochemical Studies to Establish Ternary Complex. **(A)** The left panel shows the superimposition of the crystal structures of TEAD2 in complex with flufenamic acid (PDB ID: 5DQ8) and TEAD2 in complex with palmitate (PDB ID: 5HGU). The structure of TEAD2 is shown in grey ribbon representation, while YAP is depicted in blue ribbon representation. The right panel shows a close-up view of flufenamic acid and palmitate. Amino acids within 3.5 Å of flufenamic acid are displayed in grey capped sticks representation. Amino acids from the TEAD2-palmitate complex that are within 3.5 Å of palmitate are shown in red capped sticks. Flufenamic acid is represented in capped sticks, color-coded by atom type (C, N, O, and F are shown in yellow, blue, red and light blue, respectively). Palmitate is also shown in capped sticks with carbon and oxygen atoms colored cyan and red, respectively. **(B)** Chemical structure of the bifunctional molecules designed to promote proteasomal degradation of TEADs and YAP-TAZ via cereblon. On one end (shown in pink), the bifunctional molecules are designed to bind to the TEAD palmitate pocket based on flufenamic acid analogs, while the other end (shown in blue) targets cereblon through thalidomide. Several flufenamic acid analogs are listed in the table, with both carboxylic acid and methyl ester derivatives synthesized. A cyan-colored linker was designed to tether flufenamic acid analogs to thalidomide. **(C)** A schematic representation of the HTRF biochemical assay developed to establish ternary complex formation between TEAD and cereblon in the presence of the bifunctional molecules. Antibodies fused with a d2 fluorophore and terbium cryptate were designed to bind to the His-tag on TEAD and the GST-tag on cereblon, respectively. When in proximity, the delayed emission from the terbium cryptate probe is absorbed by d2 and emitted as a fluorescence signal. Emissions from both terbium cryptate and d2 are measured to determine the extent of binding. **(D)** HTRF signal measured for the association of TEAD1 and cereblon in the presence of increasing concentrations of compounds **1** (TED-651) to **5** (TED-673). **(E)** HTRF signal measured for the association of TEAD3 and cereblon in the presence of increasing concentrations of compounds **1** (TED-651) to **5** (TED-673).

### Ternary Complex Formation

A homogeneous time-resolved fluorescence (HTRF) assay was developed to determine whether the bifunctional molecules bring TEAD and cereblon into proximity through the formation of a ternary complex (**Fig 1C**). We only tested carboxylic acid bifunctional molecules as the methyl esters require hydrolysis of ester to carboxylic acid for binding to TEAD. Recombinant His-tagged TEAD1 or TEAD3 were incubated with one of five bifunctional molecules, from **1** (TED-651) to **5** (TED-673) for 30 min, followed by the addition of GST-tagged cereblon, a GST antibody with terbium cryptate, and a His-tag antibody with a d2 fluorophore. Upon ternary complex formation, the terbium cryptate and the d2 fluorophore are brought into proximity, allowing the emission from the cryptate to excite the d2 fluorophore. For His-tagged TEAD1 (**Fig. 1D**) and His-tagged TEAD3 (**Fig. 1E**), a concentration-dependent increase in the signal intensity was observed, suggesting that the bifunctional molecules promote complex formation between TEADs and cereblon. Compounds **2** (TED-677), **3** (TED-672), and **5** (TED-673) led to the highest level of association of TEAD1 and TEAD3 with cereblon, as evidenced by a twofold increase in the HTRF signal at 10 µM. A decrease in the signal observed at higher concentrations can be attributed to the Hook effect. Compounds **1** (TED-651) and **4** (TED-675) led to substantially weaker association between TEAD1 and TEAD3 with cereblon.

To confirm that the bifunctional molecules bind to the TEAD lipid pocket, we used our HTRF assay in the presence of TED-655 (**Fig. 2A**), a small molecule that we previously developed to bind to the TEAD lipid pocket and form a covalent bond with a conserved cysteine in this pocket (39). Recombinant His-tagged TEAD3 was incubated with bifunctional molecules **2** (TED-677) [**Fig. 2B**] or **3** (TED-672) [**Fig. 2C**] for 2 and 24 h in the presence and absence of TED-655, followed by the addition of GST-tagged cereblon, a GST antibody with terbium cryptate, and a His-tag antibody with a d2 fluorophore. A time-dependent decrease in the extent of binding was observed in the presence of TED-655 for both **2** (TED-677) and **3** (TED-672). This is consistent with covalent binding of TED-655 as established previously, and it also confirms that the bifunctional molecules are engaging the TEAD lipid pocket.

**Figure 2.**
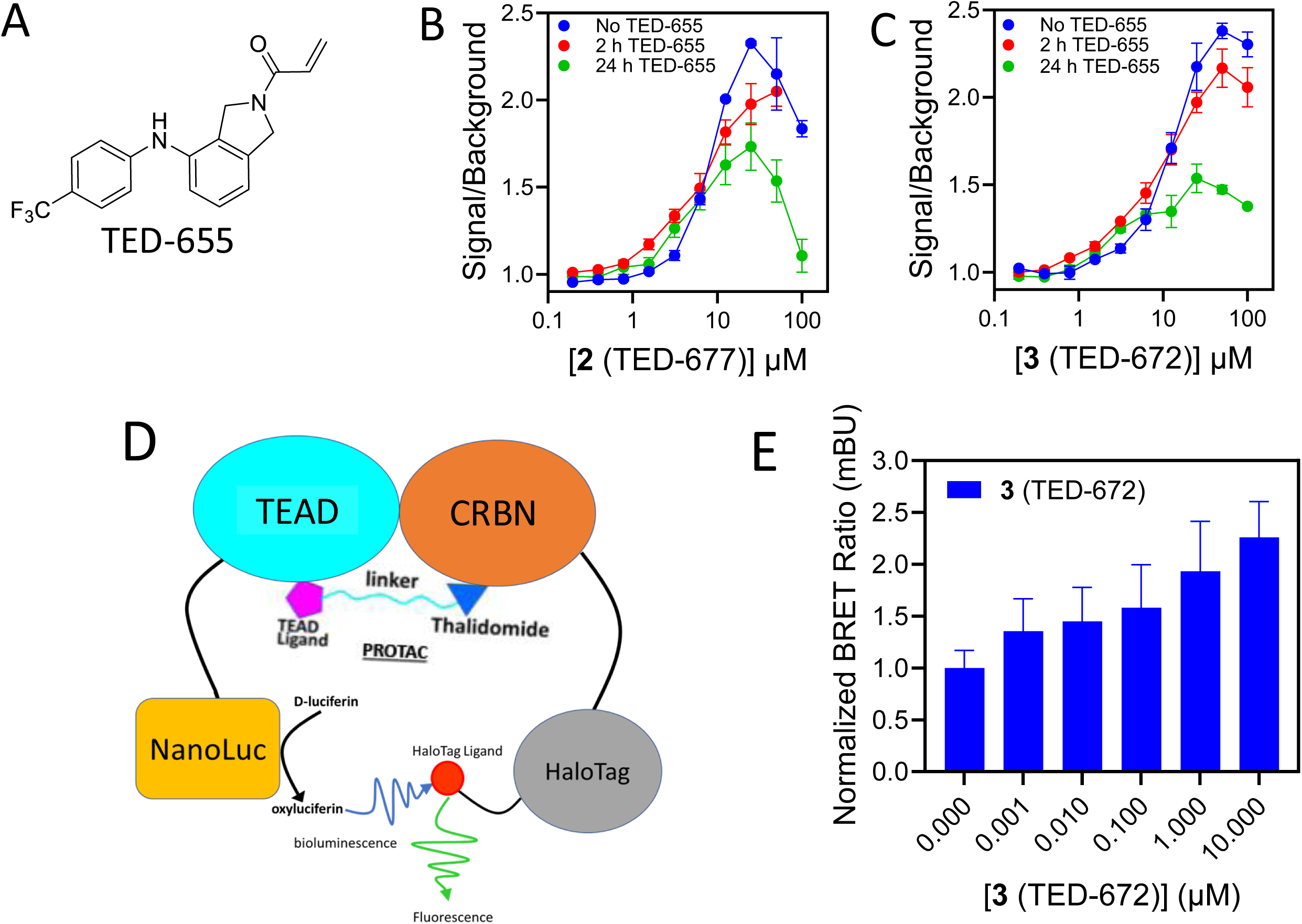
Ternary Complex Formation in Cells. **(A)** Chemical structure of TED-655. **(B)** A sample of 15 µM His-TEAD3 was incubated with 50 µM TED-655 for 2 h or 24 h at 4°C. The sample was diluted to a final concentration of 10 nM and tested for ternary complex formation to GST-CRBN with increasing concentrations of (**B**) **2 (**TED-677) or (**C**) **3 (**TED-672) as proximity inducers. (**D**) An illustration of the NanoBRET assay developed to establish ternary complex formation between bifunctional molecules, TEAD, and YAP. (**E**) HEK-293T cells were transfected with CRBN-HaloTag and TEAD1-NLuc for 24 h at a 1000:1 ratio. Cells were then treated with increasing concentrations of **3** (TED-672) for 24 h, followed by the addition of HaloTag and NLuc ligands to measure the BRET signal. These experiments were conducted in biological duplicates (mean ± s.d.; n = 6).

Next, we explored whether the bifunctional molecules promote ternary complex formation in mammalian cells. We developed a NanoBRET assay by conjugating TEAD1 with NanoLuc (TEAD1-NLuc) and cereblon with HaloTag (CRBN-Halo) (**Fig. 2D**). HEK-293T cells were transfected with TEAD1-NLuc and CBRN-Halo for 24 h, followed by treatment with increasing concentrations of **3** (TED-672) [**Fig. 2E**] for 24 h. This resulted in a concentration-dependent increase in the BRET signal. These results suggest that the heterobifunctional compounds promote ternary complex formation between TEAD and cereblon in cells.

To explore whether association resulted in reduced protein levels of TEAD, YAP, and TAZ, we resorted to western blotting analysis in MDA-MB-468 and MDA-MB-231 cell lines for **3** (TED-672) [**Fig. 3**] and its methyl ester analog **8** (TED-690) [**Fig. 4**] in biological replicates. In addition to analyzing YAP levels using a pan-YAP antibody, we also investigated the effect of the bifunctional molecules on TEAD, YAP, and TAZ protein levels. Compound **3** (TED-672) showed mild reduction in the protein levels of YAP, TEAD, TAZ, and phosphorylated YAP in a concentration-dependent manner in both MDA-MB-231 (**Fig. 3A, B**, and **C**) and MDA-MB-468 (**Fig. 3D, E**, and **F**). For example, the compound reduced YAP protein levels by 30-40% at 10 µM in MDA-MB-468 and MDA-MB-231 cells.

**Figure 3.**
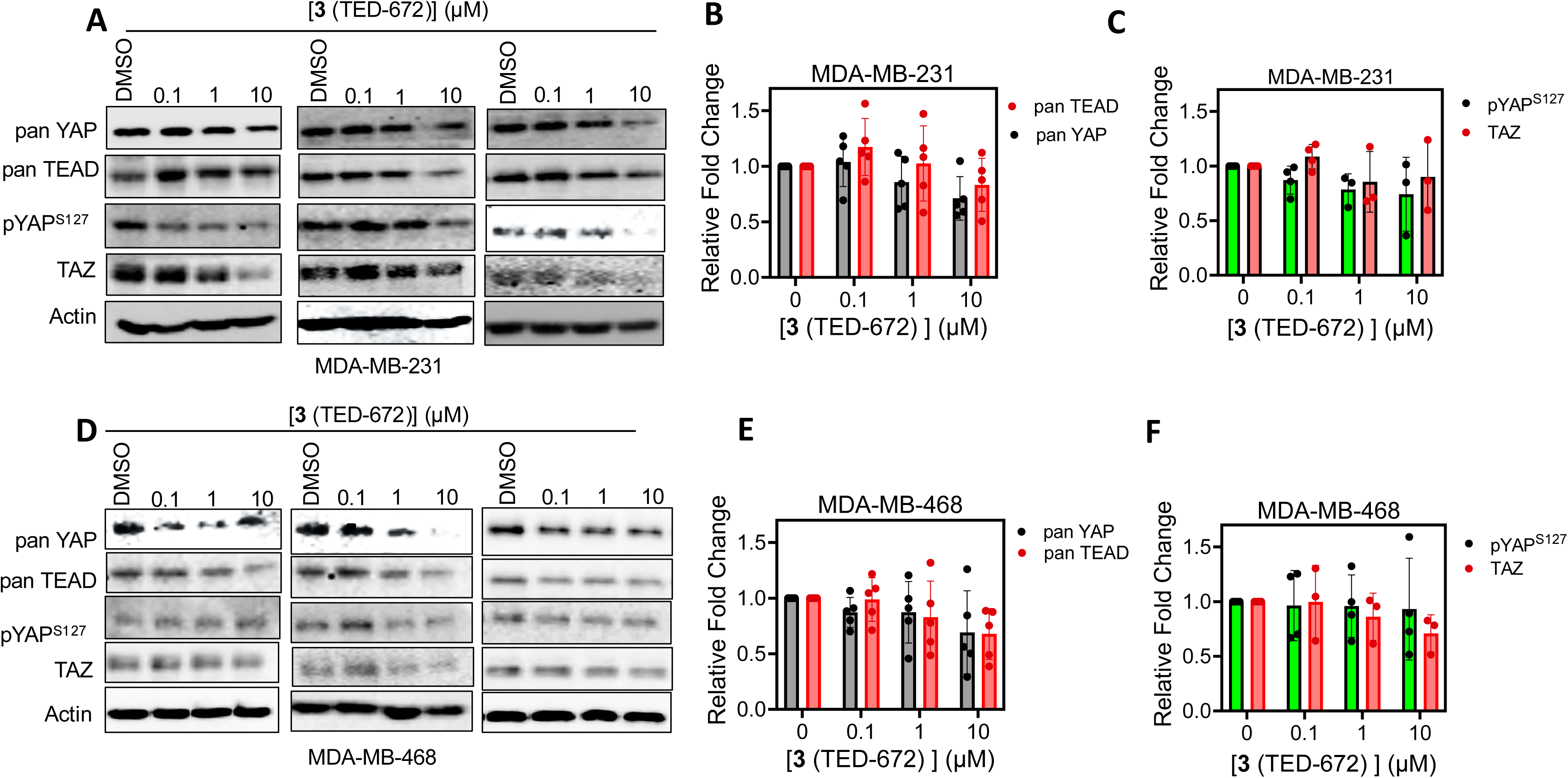
Bifunctional Carboxylic Acid Compound Tested for Protein Degradation of TEAD, YAP and TAZ. (**A**) MDA-MB-231 cells were treated with DMSO or **3** (TED-672) at three concentrations starting at 10 µM followed by 10-fold dilutions for 24 h. Pan-YAP, pYAP^S127^, pan-TEAD, TAZ and actin protein levels were detected by Western blotting in biological duplicates. Representative dose-response blots are shown. (**B**) Protein levels from the MDA-MB-231 blots were quantified using ImageJ for pan-TEAD and pan-YAP (mean ± s.d.; n = 5 replicates). **(C)** Protein levels from the MDA-MB-231 blots were quantified using ImageJ for pYAP^S127^ and TAZ (mean ± s.d.; n = 5 replicates). (**D**) MDA-MB-468 cells were treated with **3** (TED-672) at three concentrations starting at 10 µM followed by 10-fold dilutions for 24 h. Pan-YAP, pYAP^S127^, pan-TEAD, TAZ and actin protein levels were detected by Western blotting in biological duplicates. Representative dose-response blots are shown. (**E**) Protein levels from the MDA-MB-468 blots were quantified using ImageJ for pan-TEAD and pan-YAP (mean ± s.d.; n = 5 replicates). **(F)** Protein levels from the MDA-MB-468 blots were quantified using ImageJ for pYAP^S127^ and TAZ proteins (mean ± s.d.; n = 5 replicates).

**Figure 4.**
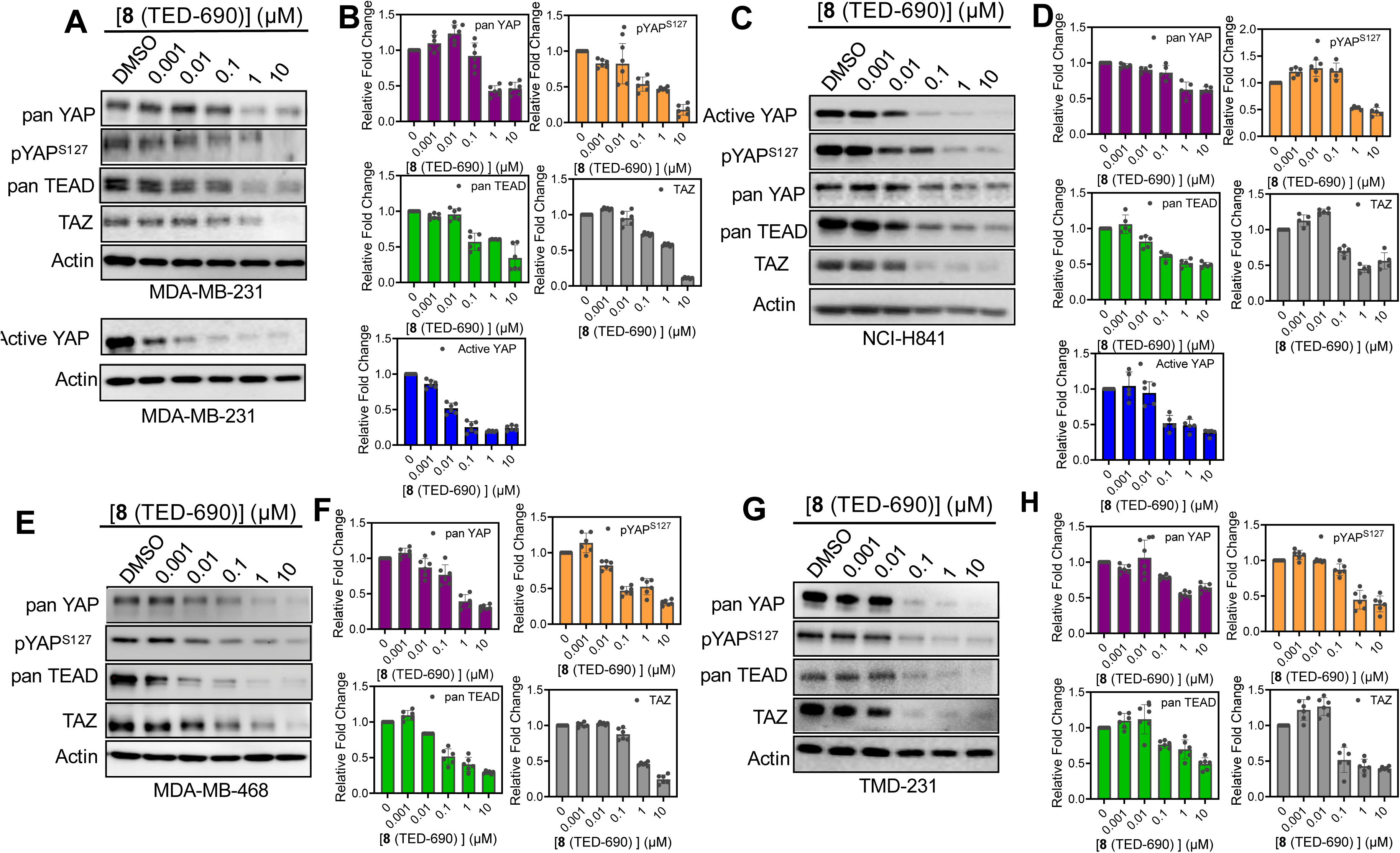
Compound 8 (TED-690) Tested for TEAD, YAP, and TAZ Protein Degradation. (**A**) MDA-MB-231 cells were treated with **8** (TED-690) at five concentrations, starting at 10 µM followed by 10-fold serial dilutions for 24 h. pan-YAP, pYAP^S127^, pan-TEAD, TAZ, active YAP, and actin protein levels were detected by Western blotting in biological replicates (mean ± s.d.; n = 6). Representative dose-response blots are shown. (**B**) Protein levels from the blots were quantified using ImageJ for pan-YAP, pYAP^S127^, pan-TEAD, TAZ, and active YAP relative to actin (mean ± s.d.; mean ± s.d.; n = 6 replicates). (**C**) NCI-H841 cells were treated with **8** (TED-690) at five concentrations, starting at 10 µM followed by serial 10-fold dilutions for 24 h. pan-YAP, pYAP^S127^, pan-TEAD, TAZ, active YAP and actin protein levels were detected by Western blotting in biological replicates (mean ± s.d.; n = 5). Representative dose-response blots are shown. **(D)** Protein levels from the blots were quantified using ImageJ for pan-YAP, pYAP^S127^, pan-TEAD, TAZ, and active YAP relative to actin (mean ± s.d.; n = 5 replicates). (**E**) MDA-MB-468 cells were treated with **8** (TED-690) at five concentrations starting at 10 µM followed by serial 10-fold dilutions for 24 h. pan-YAP, pYAP^S127^, pan-TEAD, TAZ and actin protein levels were detected by Western blotting in biological replicates (mean ± s.d.; n = 6). Representative dose-response blots are shown. (**F**) Protein levels from the blots were quantified using ImageJ for pan-YAP, pYAP^S127^, pan-TEAD and TAZ relative to actin (mean ± s.d.; n = 6 replicates). **(G)** TMD-231 cells were treated with **8** (TED-690) at five concentrations, starting at 10 µM followed by 10-fold serial dilutions for 24 h. pan-YAP, pYAP^S127^, pan-TEAD, TAZ and actin protein levels were detected by Western blotting in biological replicates (mean ± s.d.; n = 6). Representative dose-response blots are shown. (**H**) Protein levels from the blots were quantified using ImageJ for pan-YAP, pYAP^S127^, pan-TEAD and TAZ relative to actin (mean ± s.d.; n = 6 replicates).

We investigated whether methyl ester analog **8** (TED-690) exhibits improved cell permeability and leads to greater protein degradation compared to the parent carboxylic acid compound. Compound **8** (TED-690) was tested for its ability to degrade proteins in four cancer cell lines: MDA-MB-231 (**Fig. 4A** and **B**), NCI-H841 (**Fig. 4C** and **D**), MDA-MB-468 (**Fig. 4E** and **F**), and TMD-231 (**Fig. 4G** and **H**). Cells were treated for 24 h at five concentrations starting at 10 µM using 10-fold serial dilution. In MDA-MB-231 cells, the compound reduced YAP, pYAP^S127^, and active YAP protein levels with DC_50_ values of 0.1 ±0.3 µM (D_max_ = 57%), 0.1 ±0.2 µM (D_max_ = 80%), and 0.01 ±0.001 µM (D_max_ = 80%), respectively [**Table 1**]. The compound also reduced TAZ protein levels, with a DC_50_ of 0.6 ±0.3 µM (D_max_ = 89%). Finally, using a pan TEAD antibody, Western blot analysis revealed TEAD degradation with a DC_50_ of 0.1 ±0.3 µM (D_max_ = 50%). In the YAP-amplified small cell lung cancer cell line NCI-H841, **8** (TED-690) reduced YAP, pYAP^S127^, and active YAP protein levels, with DC_50_ values of 0.2 ±0.2 µM (D_max_ = 61%), 0.3 ±0.3 µM (D_max_ = 62%), and 0.03 ±0.02 µM (D_max_ = 61%), respectively (**Table 1**). The compound also led to decrease in TAZ protein levels, with a DC_50_ of 0.1 ±0.6 µM (D_max_ = 57%). Notably, TEAD protein levels were reduced with a nanomolar DC_50_ of 0.02 ±0.01 µM (D_max_ of 54%). In the triple-negative breast cancer cell line MDA-MB-468, **8** (TED-690) degraded pYAP^S127^ and TEAD with nanomolar DC_50_ values of 0.03 ±0.04 µM (D_max_ = 70%) and 0.02 ±0.03 µM (D_max_ = 71%), respectively (**Table 1**). Degradation of YAP and TAZ were observed at the sub-micromolar level, with DC_50_ values of0.1 ± 0.1 µM (D_max_ = 70%) and 0.5 ± 0.1 µM (D_max_ = 76%), respectively. Finally, in TMD-231 cells, **8** (TED-690) reduced the protein levels of YAP, pYAP^S127^, and TEAD at the sub-micromolar level, with DC_50_ values of 0.2 ± 0.1 µM (D_max_ = 46%), 0.2 ± 0.03 µM (D_max_ = 67%), and 0.2 ± 0.3 µM (D_max_ = 61%), respectively. Notably, TAZ levels were more substantially decreased in the nanomolar range (DC_50_ = 0.04 ± 0.1 µM; D_max_ = 60%).

**Table 1.** Degradation Levels Detected for Bifunctional Compounds in Cancer Cells.

|  |  | TMD-231 |  | MDA-MB-468 |  | MDA-MB-231 |  | NCI-H841 |  |
| --- | --- | --- | --- | --- | --- | --- | --- | --- | --- |
| Compound | Protein | DC<sub>50</sub> (μM) | D<sub>max</sub> (%) | DC<sub>50</sub> (μM) | D<sub>max</sub> (%) | DC<sub>50</sub> (μM) | D<sub>max</sub> (%) | DC<sub>50</sub> (μM) | D<sub>max</sub> (%) |
| 8 (TED-690) | Active YAP | - | - | - | - | 0.01 ± 0.001 | 80 | 0.03 ± 0.02 | 61 |
|  | pan YAP | 0.2 ± 0.1 | 46 | 0.1 ± 0.1 | 70 | 0.1 ± 0.3 | 57 | 0.2 ± 0.2 | 61 |
|  | pYAP<sup>S127</sup> | 0.2 ± 0.03 | 67 | 0.03 ± 0.04 | 70 | 0.1 ± 0.2 | 80 | 0.3 ± 0.3 | 62 |
|  | pan TEAD | 0.2 ± 0.3 | 61 | 0.02 ± 0.03 | 71 | 0.1 ± 0.3 | 50 | 0.02 ± 0.01 | 54 |
|  | TAZ | 0.04 ± 0.1 | 60 | 0.5 ± 0.1 | 76 | 0.6 ± 0.3 | 89 | 0.1 ± 0.6 | 57 |

The ability of **8** (TED-690) to degrade TEAD, YAP, and TAZ prompted us to explore another methyl ester, **6** (TED-650), for its potential to degrade these proteins in MDA-MB-468 and TMD-231 cell lines (**Fig. 5** and **Table 2**). Compound **6** (TED-650) reduced pan-YAP and pYAP^S127^ protein levels in the MDA-MB-468 cell line (**Fig. 5A** and **B**), with DC_50_ values of 0.07 ± 0.02 µM (D_max_ = 48%) and 0.04 ± 0.04 µM (D_max_ = 50%), respectively. In the TMD-231 cell line (**Fig. 5C** and **D**), the compound exhibited DC_50_ values of 0.3 ± 0.2 µM (D_max_ = 49%) and 0.1 ± 0.01 µM (D_max_ = 65%), respectively. Additionally, the compound reduced TEAD and TAZ protein levels in both cell lines. In TMD-231 TEAD was degraded with DC_50_ of 0.4 ± 0.1 μM (D_max_ = 63%), and TAZ with DC_50_ of 0.4 ± 0.3 μM (D_max_ = 66%). In MDA-MB-468 TEAD was degraded with DC_50_ of 0.6 ± 0.2 μM (D_max_ = 68%), and TAZ with DC_50_ of 0.4 ± 0.1 μM (D_max_ = 72%). To further investigate the species-specificity of our PROTACs, we tested **6** (TED-650) (**Fig. 5E** and **F**) and **8** (TED-690) [**Fig. 5G** and **H**] in 4T1 cells, a murine breast cancer cell line that expresses a different cereblon variant from human cells. In 4T1 cell line, neither **6** (TED-650) nor **8** (TED-690) degraded YAP degradation in contrast to their potent activity in human cell lines.

**Figure 5.**
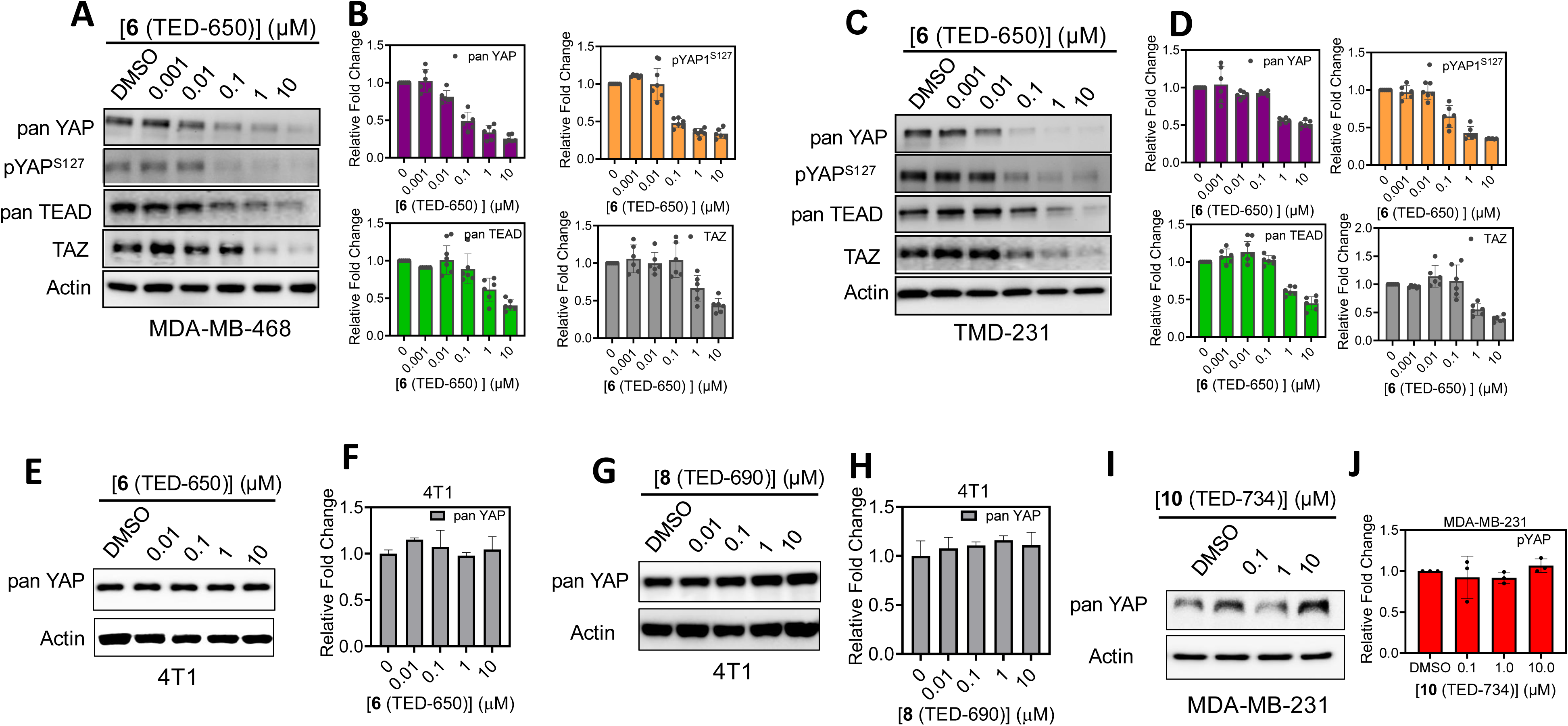
Compound 6 (TED-650) and 8 (TED-690) Tested for TEAD, YAP, and TAZ Protein Degradation. (**A**) MDA-MB-468 cells were treated with **6** (TED-650) at five concentrations starting at 10 µM followed by 10-fold serial dilutions for 24 h. pan-YAP, pYAP^S127^, pan-TEAD, TAZ, and actin protein levels were detected by Western blotting in biological replicates (mean ± s.d.; n = 6). Representative dose-response blots are shown. (**B**) Protein levels from the blots were quantified using ImageJ for pan-YAP, pYAP^S127^, pan-TEAD, and TAZ relative to actin (mean ± s.d.; n = 6 replicates). (**C**) TMD-231 cells were treated with **6** (TED-650) at five concentrations starting at 10 µM followed by 10-fold serial dilutions for 24 h. pan-YAP, pYAP^S127^, pan-TEAD, TAZ and actin protein levels were detected by Western blotting in biological replicates (mean ± s.d.; n = 6). Representative dose-response blots are shown. **(D)** Protein levels from the blots were quantified using ImageJ for YAP, pYAP^S127^, pan-TEAD and TAZ relative to actin (mean ± s.d.; n = 6 replicates). (**E**) 4T1 cells were treated with **6** (TED-650) at four concentrations starting at 10 µM followed by 10-fold serial dilutions for 24 h. pan-YAP and actin protein levels were detected by Western blotting in biological replicates (mean ± s.d.; n = 3). Representative dose-response blots are shown. (**F**) Protein levels from the blots were quantified using ImageJ for pan-YAP and relative to actin (mean ± s.d.; n = 3 replicates). (**G**) 4T1 cells were treated with **8** (TED-690) at four concentrations starting at 10 µM followed by 10-fold serial dilutions for 24 h. pan-YAP and actin protein levels were detected by Western blotting in biological replicates (mean ± s.d.; n = 3). Representative dose-response blots are shown. **(H)** Protein levels from the blots were quantified using ImageJ for pan-YAP and relative to actin (mean ± s.d.; n = 3 replicates). (**I**) MDA-MB-231 cells were treated with **10** (TED-734) at three concentrations starting at 10 µM followed by 10-fold serial dilutions for 24 h. pan-YAP and actin protein levels were detected by Western blotting in biological replicates (mean ± s.d.; n = 3). Representative dose-response blots are shown. **(J)** Protein levels from the blots were quantified using ImageJ for pan-YAP and relative to actin (mean ± s.d.; n = 3 replicates).

**Table 2.** Degradation Levels Detected for Bifunctional Compounds in Cancer Cells.

|  |  | TMD-231 |  | MDA-MB-468 |  |
| --- | --- | --- | --- | --- | --- |
| Compound | Protein | DC<sub>50</sub> (μM) | D<sub>max</sub> (%) | DC<sub>50</sub> (μM) | D<sub>max</sub> (%) |
| 6 (TED-650) | pan YAP | 0.3 ± 0.2 | 49 | 0.02 ± 2.7 | 48 |
|  | pYAP<sup>S127</sup> | 0.1 ± 0.01 | 65 | 0.04 ± 0.04 | 50 |
|  | pan TEAD | 0.4 ± 0.1 | 63 | 0.6 ± 0.2 | 68 |
|  | TAZ | 0.4 ± 0.3 | 66 | 0.4 ± 0.1 | 72 |

Additionally, we tested **10** (TED-734) in the human breast cancer cell line MDA-MB-231 to establish that degradation was mediated by cereblon (**Fig. 5I** and **J**). Compound **10** (TED-734) contains a methylated thalidomide moiety, which prevents thalidomide from binding to cereblon, and therefore serves as a negative control. As expected, we did not observe degradation of pan-YAP, which further supports that the bifunctional molecules degrade proteins in a cereblon-dependent manner.

### Time-Dependent Effect of Bifunctional Compounds

To gain insight into the time-dependent effects on protein levels, we examined the degradation of YAP, TEAD and TAZ at different time points (**Fig. 6**). These studies were conducted using **8** (TED-690) in MDA-MB-231 and NCI-H841 cells. In each case, cells were incubated with compound at 1 µM for 2, 8, 16, and 24 h. Compound **8** (TED-690) demonstrated time-dependent degradation, reducing YAP, TEAD, and TAZ protein levels in MDA-MB-231 cells with degradation half-lives of 10.1 ± 1.9 h, 9.6 ± 1.4 h, and 9.7 ± 1.4 h, respectively (**Fig. 6A**, **B**, and **E**). The degradation occurred more rapidly in the NCI-H841 lung cancer cell line, with half-lives of 7.3 ± 0.7 h, 7.8 ± 0.7 h, and 5.8 ± 0.2 h, respectively (**Fig. 6C, D**, and **E**).

**Figure 6.**
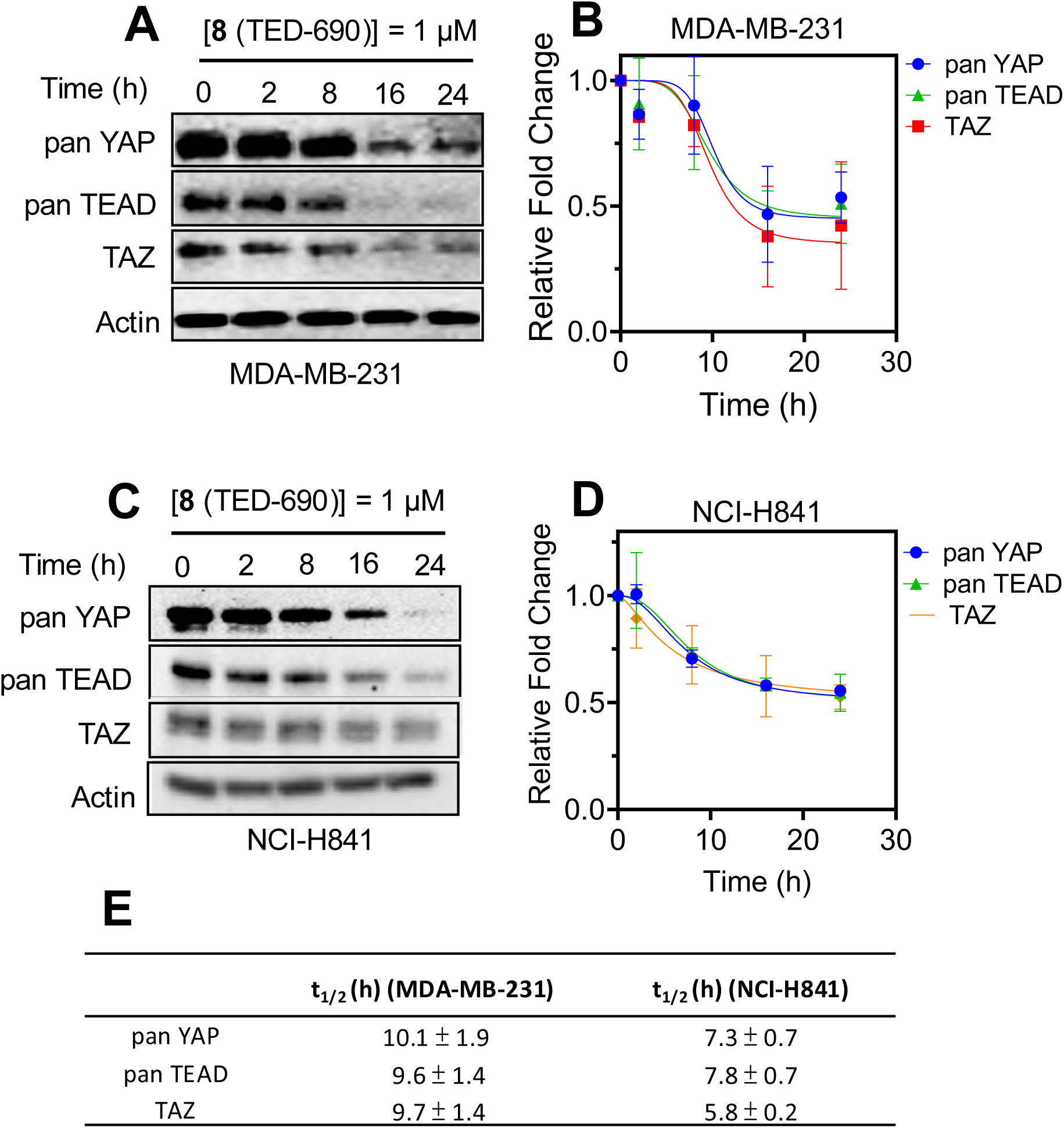
Time-Dependent Degradation of TEAD, YAP, and TAZ. (**A**) Representative blot depicting the effect of the bifunctional molecule at different time points. MDA-MB-231 cells were treated with **8** (TED-690) for 2, 8, 16, and 24 h, followed by Western blot to detect protein levels of pan-TEAD, pan-YAP, and TAZ. (**B**) Protein levels from the blots were quantified using ImageJ for pan-YAP, pan-TEAD and TAZ relative to actin (mean ± s.d.; n = 4 replicates). **(C)** Representative blot depicting the effect of the bifunctional molecule at different time points. NCI-H841 cells were treated with **8** (TED-690) for 2, 8, 16, and 24 h, followed by Western blot to detect protein levels of pan-TEAD, pan-YAP, and TAZ. (**D**) Protein levels from the blots were quantified using ImageJ for pan-YAP, pan-TEAD and TAZ relative to actin (mean ± s.d.; n = 4 replicates). (**E**) Degradation half-lives were determined for the bifunctional compound using the time-dependent Western blot studies.

### Degradation of TEAD, YAP, and TAZ Is Proteasomal

To confirm that the bifunctional molecules induce degradation through the proteasome, Western blot analyses were carried out using the proteasome inhibitors MG132 and epoxomicin (EPO) (**Fig. 7**). MDA-MB-231 cells were treated with **8** (TED-690) at 1 µM for 24 h, followed by Western blot analysis. Proteasome inhibitors were added six hours before cell harvesting (**Fig. 7A** and **B**). Treatment with the compound reduced YAP protein levels. When cells were treated with the proteasome inhibitor MG132 or EPO at 5 µM, YAP protein levels were similar to DMSO control levels. When cells were treated with both **8** (TED-690) and proteasome inhibitors, substantially less degradation was observed, suggesting that the degradation observed throughout this study is likely proteasomal. Notably, STAT3 protein levels remained unaffected by **8** (TED-690) treatment. STAT3, a protein not expected to interact with TEAD or cereblon, served as a specificity control.

**Figure 7.**
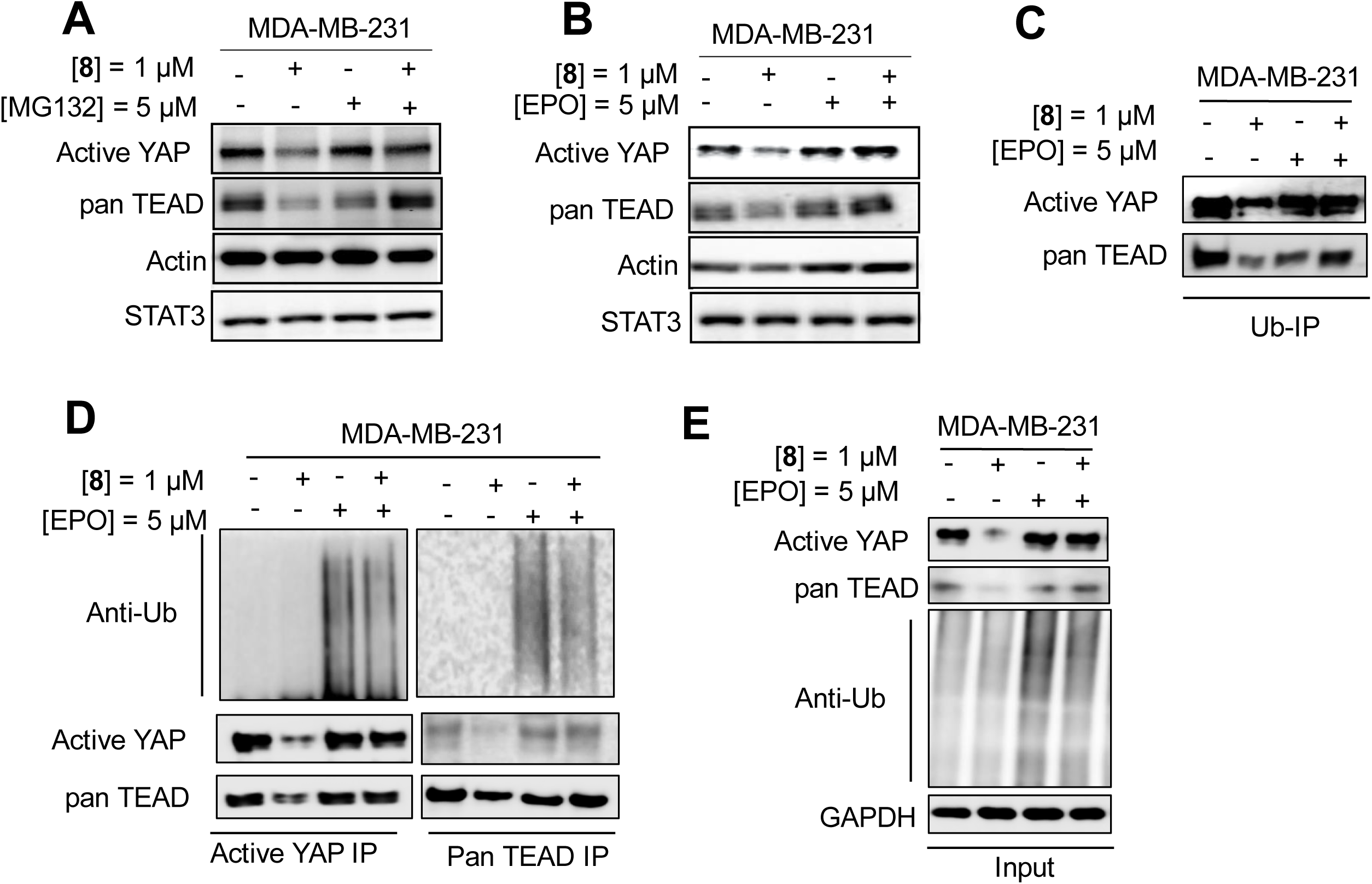
Evidence for Proteasomal Degradation of TEAD, YAP, and TAZ by Bifunctional Molecules. (**A**) MDA-MB-231 cells were treated with either 1 µM of **8** (TED-690) for 24 h, or 5 µM of MG132 for 6 h prior to harvesting the cells, or a combination of both compounds. Protein levels of active YAP, pan-TEAD, actin, and STAT3 were detected by immunoblot analysis and compared to the DMSO control. (**B**) MDA-MB-231 cells were treated with 1 µM of **8** (TED-690) for 24 h, or 5 µM of EPO for 6 h prior to harvesting, or a combination of both compounds. Protein levels of active YAP, pan-TEAD, actin, and STAT3 were detected by immunoblot analysis and compared to the DMSO control. (**C**) MDA-MB-231 were treated with **8** (TED-690), EPO, either alone or in combination, followed by Western blot analysis of active YAP and pan-TEAD levels (input control), or immunoprecipitation using antibodies against active YAP or pan-TEAD. Immunoprecipitated samples were analyzed by Western blot using an anti-ubiquitin antibody. **(D)** MDA-MB-231 cells were treated with 1 µM of **8** (TED-690) in the presence or absence of 5 µM EPO, followed by immunoprecipitation with an anti-ubiquitin antibody and subsequent Western analysis to detect protein levels of YAP and pan-TEAD. (**E**) MDA-MB-231 lysate from (D) were collected as Input, followed by Western blot analysis of active YAP and pan-TEAD levels (input control) using antibodies against active YAP, pan-TEAD, ubiquitin and GAPDH.

To further support this conclusion, we performed a co-immunoprecipitation experiment by pulling down ubiquitin, followed by Western blot analysis of YAP and TEAD protein levels (**Fig. 7C**). MDA-MB-231 cells were treated with **8** (TED-690) either alone or in combination with the proteasome inhibitor EPO. Proteins immunoprecipitated with a ubiquitin antibody were analyzed by Western blotting for active YAP and pan-TEAD protein. When cells were treated with **8** (TED-690) alone, pan-TEAD and active YAP levels remained unchanged compared to the DMSO control. However, treatment with EPO led to an increase in pan-TEAD and active YAP levels, consistent with accumulation of ubiquitinated proteins due to proteasome inhibition. Notably, in the presence of both **8** (TED-690) and EPO, an even greater increase in pan-TEAD and active YAP protein levels was observed, further supporting a proteasomal degradation mechanism.

A co-immunoprecipitation experiment was conducted to determine whether YAP and TEAD undergo ubiquitination following treatment with compound (**Fig. 7D** and **E**). MDA-MB-231 cells were treated with 1 µM **8** (TED-690) for 24 h in the presence or absence of proteasome inhibitor EPO. After treatment, cells were harvested and lysed, and lysates were incubated with beads conjugated to YAP or TEAD antibodies. Western blot analysis was then performed on both lysates (input control) and immunoprecipitated samples to detect YAP and TEAD. A substantial accumulation of ubiquitinated YAP and TEAD was observed following pull-down with either YAP or TEAD antibodies. In a repeat experiment, cells were treated with **8** (TED-690) for 24 h, followed by the addition of EPO (5 µM) six hours before harvesting. No effect was detected for YAP and TEAD levels in the input control sample. However, ubiquitination was detected on both YAP and TEAD, consistent with proteasomal degradation.

### Bifunctional Molecules Inhibit Transcriptional Activity and Cell Viability

We explored the effects of **8** (TED-690) on TEAD transcriptional activity using qRT-PCR analysis in MDA-MB-231, MDA-MB-468, TMD-231, and NCI-H841 cells (**Fig. 8A** and **B**). The compound reduced CTGF mRNA levels in a concentration-dependent manner across all four cell lines. The effect was most pronounced in TMD-231 and NCI-H841 cells, with IC_50_ values of 0.1 ± 0.05 µM and 0.1 ± 0.02 µM, respectively. In contrast, reductions in CTGF mRNA levels relative to the DMSO control occurred at higher IC_50_ values in the micromolar range for MDA-MB-231 (1.0 ± 0.6 µM) and MDA-MB-468 (2.4 ± 0.7 µM).

**Figure 8.**
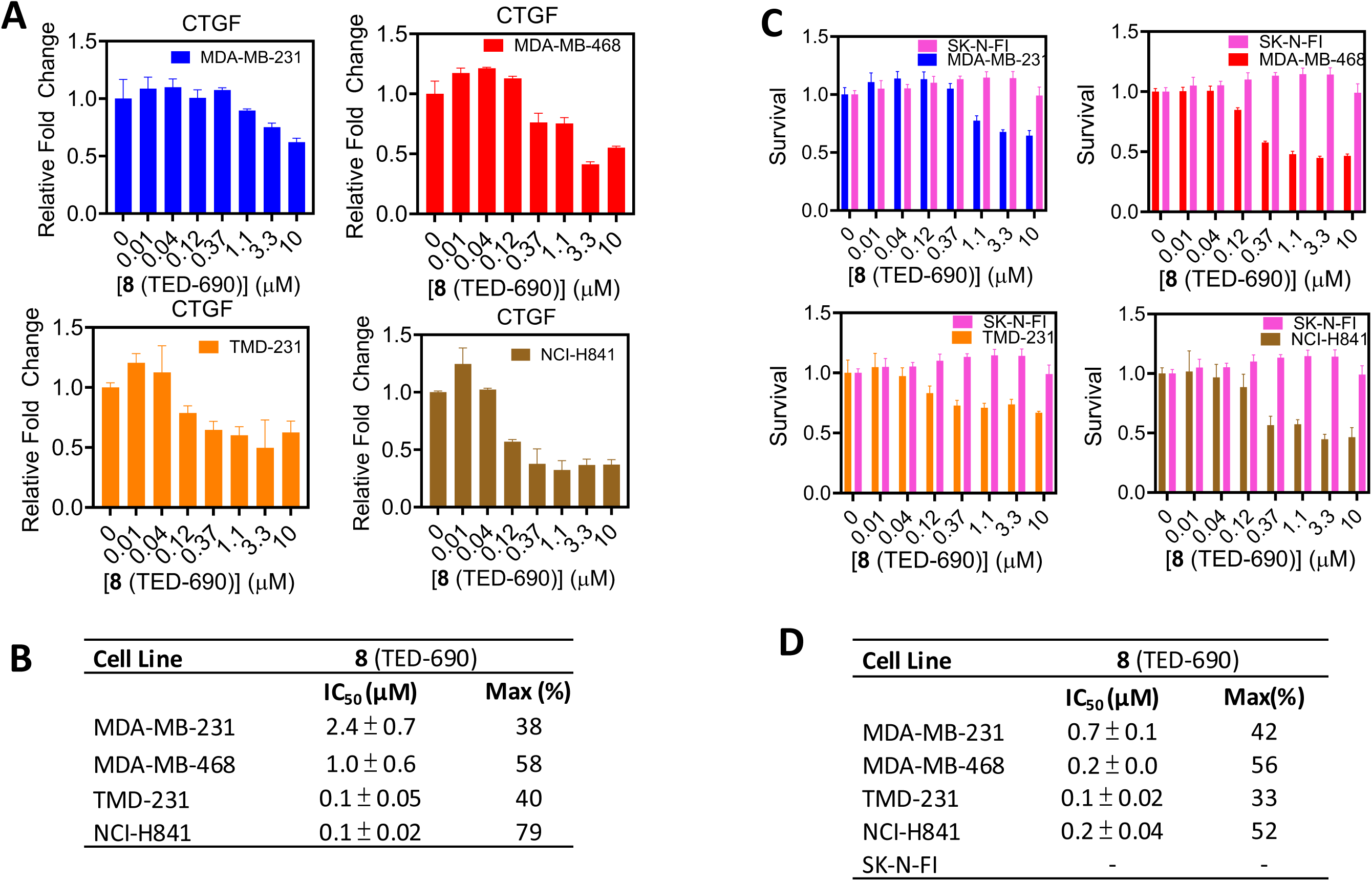
PROTAC YAP Degraders Inhibit YAP-TEAD Transcriptional Activity and Cell Survival. (**A**) MDA-MB-468, TMD-231, MDA-MB-231 and NCI-H841 cells were treated with **8** (TED-690), starting at 10 µM followed by 3-fold serial dilutions, and analyzed using qRT-PCR. (**B**) Inhibition curves were generated to determine IC_50_ (µM) values for the inhibition of *CTGF* mRNA levels by the compound. (**C**) MDA-MB-468, TMD-231, NCI-H841, MDA-MB-231 and SK-N-FI cells were cultured in monolayer format and treated with **8** (TED-690) for 3 days at concentrations starting at 10 µM, followed by serial dilutions. (**D**) Inhibition curves were generated to determine IC_50_ (µM) for the inhibition of cell viability (mean ± s.d., n = 3 biological replicates).

We investigated the effects of **8** (TED-690) on the cell viability of MDA-MB-231, MDA-MB-468, TMD-231, NCI-H841, and SK-N-FI cells (**Fig. 8C** and **D**). SK-N-FI cells do not express YAP and, therefore, should not be affected by the compound. Cells were treated with **8** (TED-690) for three days, followed by quantification of ATP levels using the Promega CellTiter-Glo assay as a measure of cell viability. The compound inhibited the viability of breast and lung cancer cell lines, with IC_50_ values in the sub-micromolar range. As expected, the compound had no effect on SK-N-FI cell viability.

## DISCUSSION

Interest in the Hippo pathway and its components has grown substantially over the past decade. The roles of the Hippo pathway transcription factor TEAD and its coactivators YAP and TAZ in cancer are well established. Upon binding to TEAD, YAP or TAZ initiates the transcription of target genes associated with tumorigenesis, growth and metastasis. The discovery that TEADs contain a palmitate-binding pocket located outside the TEAD•YAP-TAZ interface has led to the development of a large number of small molecules targeting this pocket, several of which have entered clinical trials. Although these compounds do not affect TEAD binding to YAP, they inhibit TEAD transcriptional activity. More recently, Novartis developed small molecules that target TEAD at the TEAD•YAP-TAZ protein-protein interface, and these compounds have also reached clinical trials. In addition to inhibitors, proteolysis-targeted chimeras (PROTACs) have been developed for the degradation of TEAD.

The TEAD transcription factor is unique in that it lacks a transactivation domain, which is instead located on YAP and TAZ. This characteristic allows YAP and TAZ to initiate transcriptional activity independently of TEADs. YAP and TAZ have also been shown to interact with other transcription factors, such as AP-1 and RUNX, to drive TEAD-independent transcriptional activity. Additionally, YAP and TAZ are known to bind chromatin for epigenetic regulation independently of TEADs. Consequently, small molecules that bind to or degrade TEADs are unlikely to affect TEAD-independent YAP and TAZ activities.

We wondered whether a small molecule that binds to TEADs and possesses a sufficiently long linker with a ubiquitin ligase ligand tethered at its other end could facilitate the ubiquitination and degradation of YAP and TAZ. To explore this possibility, we designed a bifunctional molecule based on flufenamic acid, a small molecule previously shown to bind to the palmitate pocket of TEADs. We modified the compound with substituents that mimic the binding pose of palmitate and enhance its affinity. Additionally, we designed a linker extending beyond the palmitate pocket to enable ubiquitin ligase recruitment, facilitating the covalent attachment of ubiquitin to YAP or TAZ, leading to their degradation.

Flufenamic acid contains a carboxylic acid, which is also present in the five derivatives we designed. Since bifunctional molecules must cross two membranes, we suspected that the carboxylic acid might hinder their ability to reach the transcription factor in the nucleus. To address this, we synthesized methyl ester derivatives for each of the five compounds. We then conducted an initial screen using Western blot analysis with a YAP antibody to identify bifunctional molecules that degrade YAP. Several of these compounds were found to induce YAP degradation in a concentration-dependent manner in MDA-MB-468 cells.

Follow-up studies assessing not only YAP but also TEAD and TAZ revealed that the methyl esters induced more profound degradation than the corresponding carboxylic acids. Initially, we selected carboxylic acid **3** (TED-672) and corresponding methyl ester counterpart **8** (TED-690) for comparison. The carboxylic acid degraded with DC_50_ values of approximately 10 µM, whereas the methyl ester demonstrated significantly greater potency, with sub-micromolar and nanomolar DC_50_ values. Further time-dependent studies showed that the degradation half-life of the ester compounds was nearly 10 hours. This finding is consistent with a prodrug mechanism, in which the conversion of the ester to the active carboxylic acid form within cells may require several hours to occur.

We tested the effect of the compounds on active nuclear YAP levels and phosphorylated YAP. Interestingly, the compounds degraded both active and phosphorylated YAP. This was unexpected, as the standard model for Hippo signaling suggests that phosphorylated YAP is either sequestered in the cytoplasm by 14-3-3 proteins or degraded in the proteasome. However, some studies have indicated that phosphorylated YAP can also translocate to the nucleus (40, 41). Our results suggest that phosphorylated YAP is not only present in the nucleus but may also form a complex with TEADs. Whether this complex contributes to transcriptional activity remains unclear. The observation that our compounds are significantly more effective at degrading active YAP compared to phosphorylated YAP may be due to phosphorylation-induced changes in YAP positioning relative to cereblon, potentially influencing ubiquitination and degradation.

In this work, we show that heterobifunctional molecules that bind to the TEAD lipid pocket have the potential to degrade not only TEAD, but also its binding partners YAP and TAZ. In one mechanism, TEAD is ubiquitinated and transported along with YAP and TAZ to the proteasome where the complex is degraded. In another mechanism, YAP and TAZ are directly ubiquitinated while in complex with TEAD. This is possible since the bifunctional molecules bind to an allosteric pocket on TEAD close to the TEAD•YAP/TAZ interface. It is expected that thalidomide or analogs will recruit cereblon close to both TEAD and YAP/TAZ enabling formation of a ternary complex with either TEAD or YAP/TAZ for ubiquitination and degradation. Future studies will establish the mechanism of degradation and will explore whether TEAD-independent YAP and TAZ transcriptional activities are also affected.

## METHOD DETAILS

### Recombinant Protein Expression and Purification

HIS-TEAD1 (residues 209-426) was cloned into pET-28a-TEV vector and transformed into *E. coli* BL21 (DE3) cells for expression. The cells were grown at 37°C to an OD_600_ of 0.6 and induced with 0.5 mM isopropyl β-D-1-thiogalactopyranoside (IPTG) overnight at 16 °C. Cells were harvested by centrifugation and lysed using a microfluidizer in lysis buffer [2 x phosphate buffered saline (PBS), 2 mM β-mercaptoethanol (BME)]. The lysate was clarified by centrifugation and the supernatant was loaded onto a 5-mL HisTrap HP column (Cytiva Life Sciences, Marlborough, MA). The column was washed with 10 column volume (CV) of lysis buffer and eluted using a linear gradient against elution buffer (2 x PBS, 500 mM imidazole, pH 7.8, 2 mM BME). The eluted protein was concentrated and treated with 6 mM hydroxylamine for 2.5 h at room temperature (RT). The sample was further purified using a Superdex 75 pg XK16/60 size-exclusion chromatography (SEC) column (Cytiva Life Sciences, Marlborough, MA) in PBS containing 1 mM dithiothreitol (DTT).

HIS-TEAD3 (residues 216-435) was cloned into the pET-28a-TEV vector and transformed into *E. coli* BL21 (DE3) cells for expression. The cells were grown at 37°C to an OD_600_ of 0.6 and induced with 0.5 mM IPTG overnight at 16°C. Cells were harvested by centrifugation and lysed using a microfluidizer in lysis buffer (500 mM NaCl, 20 mM NaPi pH 7.8, 2 mM BME). The lysate was clarified by centrifugation and the supernatant was loaded onto a 5-mL HisTrap HP column (Cytiva Life Sciences, Marlborough, MA). The column was washed with 10 CV of lysis buffer and eluted using a linear gradient against elution buffer (1 M NaCl, 0.5 M imidazole, 2 mM BME, pH 6.5). The eluent was concentrated and treated with 6 mM hydroxylamine for 2.5 h at RT. The sample was further purified using a Superdex 75pg XK16/60 SEC column (Cytiva Life Sciences, Marlborough, MA) in PBS containing 2 mM DTT.

GST-CRBN (residues 318-426) was cloned into the pGEX-6P-3 vector and transformed into *E. coli* BL21 (DE3) cells. The cells were grown in lysogeny broth (LB) at 37°C to an OD_600_ of 0.6 and induced with 0.5 mM IPTG overnight at 18°C. Cells were harvested by centrifugation and lysed using a microfluidizer in lysis buffer (20 mM Tris pH 8.0, 500 mM NaCl, 1 mM tris(2-carboxyethyl) phosphine (TCEP)). The lysate was clarified by centrifugation and the supernatant was loaded onto a 5-mL GSTrap HP column (Cytiva Life Sciences, Marlborough, MA). The column was washed with 12 CV of lysis buffer and eluted with 5 CV of elution buffer (10 mM glutathione, 20 mM Tris pH 8.0, 500 mM NaCl, 1 mM TCEP). The eluent was further purified using a Superdex 200pg XK26/60 SEC column ((Cytiva Life Sciences, Marlborough, MA) in 50 mM HEPES, pH 7.4, 200 mM NaCl, and 1 mM TCEP.

### Homogeneous Time-Resolved Fluorescence (HTRF) Assay

In a white 384-well ProxiPlate (Catalog Number 6008280, Perkin Elmer, Waltham, MA) 5 µL of 40 nM His-tagged TEAD1 or TEAD3 in assay buffer (PBS, 2 mM DTT) was aliquoted. Serially diluted compounds (0.2-100 µM final concentration) in 5 µL of 4% (v/v) dimethyl sulfoxide (DMSO) were added and incubated for 30 min at room temperature. After the incubation, 5 µL of 40 nM GST-CRBN was added and incubated for an additional 30 min at room temperature. An Anti-His antibody fluorescently tagged with d2 (Catalog Number 61HISDLA, Revvity, Waltham, MA) and anti-GST antibody fluorescently tagged with terbium cryptate were added sequentially with a 30 min interval between additions, to a final concentration of 1 µg per well for each antibody. The plate was incubated for 60 min at room temperature and read on an Envision Multiplate Reader (Perkin Elmer, Waltham, MA) using an excitation filter at 320 nm and emission filters at 615 nm and 665 nm. The time delay for the measurement was set to 120 µs.

### Western blotting

MDA-MB-231, TMD-231, MDA-MB-468 and NCI-H841 were seeded in 24-well plates at a density of 5 x 10^4^ cells per well in complete media. Cells were treated with the indicated compounds for 24 h at specified concentrations. For time-course studies, MDA-MB-231 and NCI-H841 cells were incubated with the compounds at the indicated time points and harvested for Western blot analysis. For proteasome inhibitor studies, MDA-MB-231 cells were treated with DMSO or the indicated compounds for 24 h. Cells were then treated with MG132 (VWR, PA) or epoxomicin (EPO) (Sellechem, TX) at 5 µM for 6 hours prior to harvesting. Total cell lysates were collected in lysis buffer (Cell Signaling, MA) supplemented with protease and phosphatase inhibitors (ThermoFisher Scientific, MA) and quantified using the BCA assay (VWR, PA). Proteins were separated by SDS-PAGE and transferred to PVDF membrane (Bio-Rad, CA). Membranes were incubated with the indicated primary antibodies overnight at 4°C, followed by incubation with HRP-linked anti-rabbit or anti-mouse secondary antibodies for signal detection.

### Immunoprecipitation

MDA-MB-231 cells were seeded in 10 cm plates with complete medium. Cells were treated with compounds at indicated concentrations for 24 h at low cell density. Subsequently, cells were exposed to the proteasome inhibitor epoxomicin (EPO) (Sellechem, TX) at 5 µM for 6 hours prior to harvesting. Total cell lysates were prepared using lysis buffer (Cell Signaling, MA) supplemented with protease and phosphatase inhibitors (ThermoFisher Scientific, MA) and quantified by the BCA assay (VWR, PA). Proteins were immunoprecipitated with the specified antibodies, separated by SDS-PAGE, and transferred to PVDF membranes (Bio-Rad, CA). Membranes were incubated with the indicated primary antibodies overnight at 4°C, followed by probing with HRP-linked anti-rabbit or anti-mouse secondary antibodies.

### NanoBRET Assay

HEK-293T cells were plated in 6-well plates at 3 x 10^5^ cells/well prior to transfection. After overnight incubation, the cells were co-transfected with plasmids encoding CRBN tagged with HaloTag (CRBN-Halo) at the C-terminus and TEAD1 tagged with NanoLuc (TEAD1-NLuc) at the C-terminus using Lipofectamine 3000 (Invitrogen, Waltham, MA). A donor-to-acceptor (TEAD-NLuc and CRBN-Halo) transfection ratio of 1000:1 was used. After 24 h post-transfection, cells were re-seeded in white, opaque 96-well plates with 2 x 10^4^ cells/well in Opti-MEM I Reduced Serum Medium (ThermoFisher Scientific, MA) supplemented with 4% FBS and 100 nM HaloTag ligand, with and without the 618 ligand. Cells treated with DMSO or compounds were incubated for additional 24 h at 37 °C. After treatment, NanoBRET signals were measured following the addition of NanoBRET furimazine substrate at 10 µM, and BRET signals were recorded within 10 min using a Biotek Synergy Neo2 Plate Reader (λ_Emission_ = 460 ± 40 nm and λ_Emission_ = 618 ± 40 nm). NanoBRET ratios (milliBRET units; mBU) were calculated as follow: BRET = 618 nm/460 nm × 1000. Corrected NanoBRET ratios were obtained by subtracting the DMSO control values.

### RNA Extraction and Real-Time PCR (Polymerase Chain Reaction)

MDA-MB-468, MDA-MB-231, TMD-231 and NCI-H841 cells were incubated with DMSO or the indicated compounds in a dose-dependent manner for 24 h. Cellular RNA was extracted and converted to cDNA following the vendor’s protocol (Bio-Rad, Hercules, CA). YAP/TEAD1-4 mediated transcriptional activation was assessed using *CTGF*/*GAPDH* as readouts. The sequences of primers are listed in the **Star Table**.

### Cell Viability Assay

MDA-MB-468, MDA-MB-231, TMD-231, NCI-H841 and SK-N-FI cells were seeded in 96-well plates at a density of 10^3^ cells/well in 100 µL medium for 24 h at 37°C. Subsequently, the cells were exposed to 100 µL of medium containing DMSO or the indicated compounds. Cultures were further incubated at 37°C for 3 days. Measurements were performed according to the manufacturer’s instructions (Promega, Madison, WI). In brief, 100 µL of CellTiter-Glo 2.0 reagent was added directly to the wells, followed by centrifugation for 1 minute. Cells were then incubated at room temperature for 10 minutes in the dark. Luminescence signals were measured, and background absorbance was corrected using wells containing medium only (no cells).

## STAR METHODS

### KEY RESOURCES TABLE

#### STAR METHOD

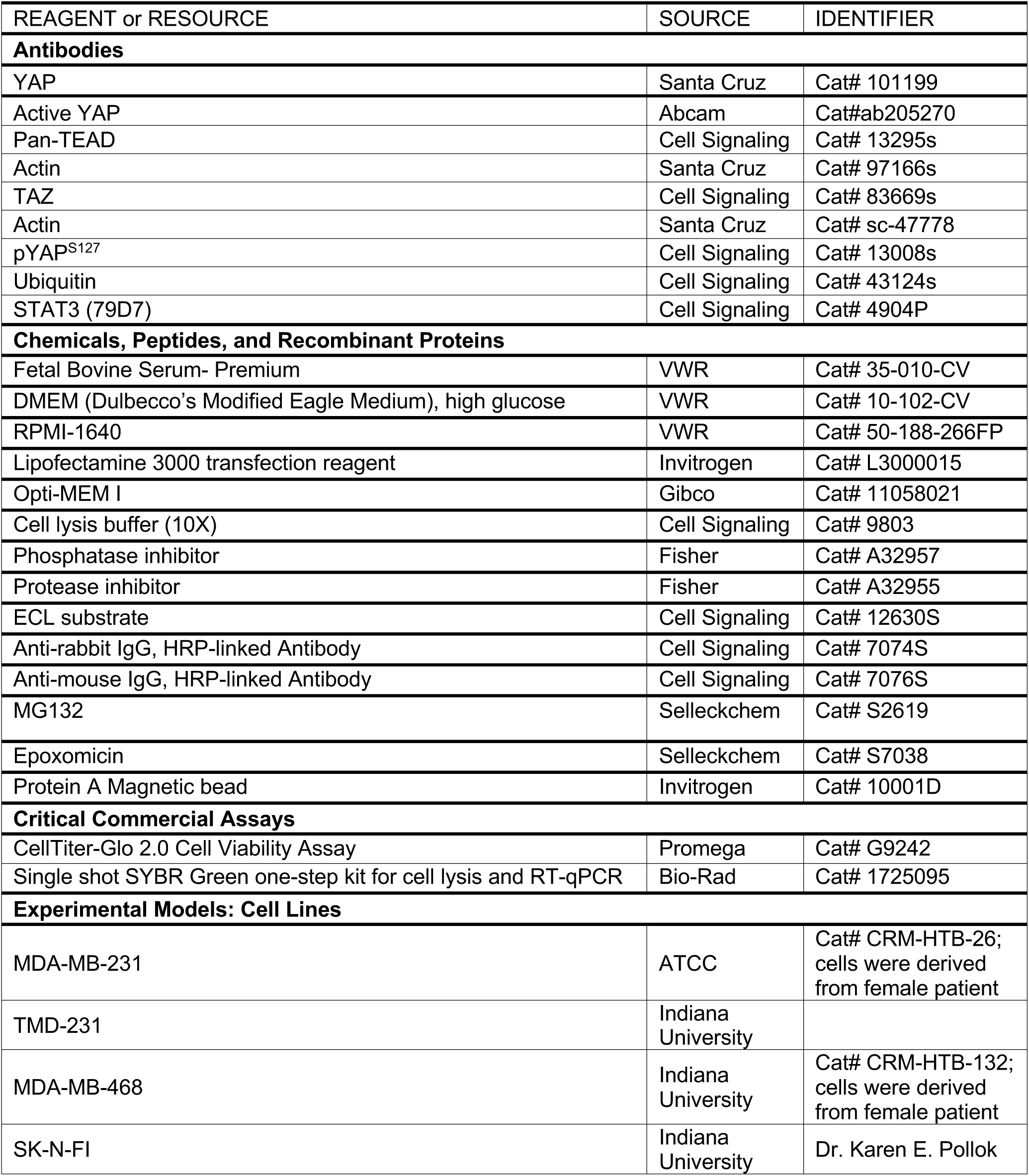
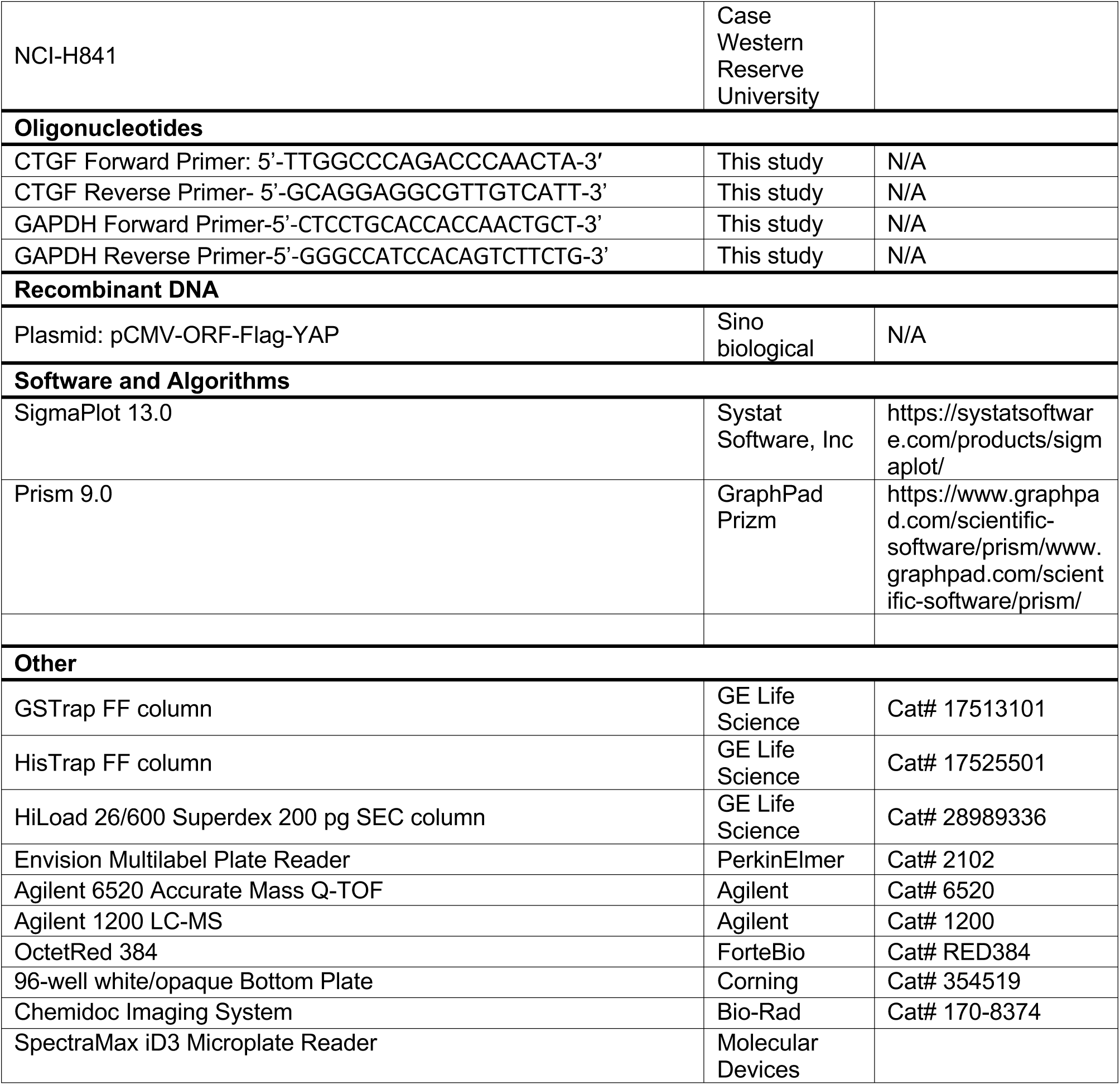

## CONTACT FOR REAGENT AND RESOURCE SHARING

Further information and requests for resources and reagents should be directed to, and will be fulfilled by, the Lead Contact, Samy O. Meroueh.

## SUPPORTING INFORMATION

Analytical data for heterobifunctional compounds.

## AUTHOR CONTRIBUTIONS

I.-J.Y. and M.K.G carried out all biological studies. K.B.-E. and M.K.G. carried out biochemical studies. S.O.M., I.-J.Y., and K.B.-E. designed experiments and analyzed data. S.O.M. designed all compounds. S.O.M., I.-J.Y., and K.B.-E. wrote the paper.

## ACKNOWLEDGMENTS

The research was supported by an American Cancer Society Research Scholar Grant RSG-12-092-01-CDD (SOM), a Vera Bradley Foundation fellowship (KB), a Vera Bradley Foundation grant (SOM), an Indiana University Simon Cancer Center Near Miss Initiative grant (SOM), and the 100 Voices of Hope (SOM). The research was supported by the Department of Veteran Affairs (I01BX005188) [SOM], and the National Institutes of Health (R01CA264471) [SOM].

