## Supplemental Information for "Degradation of the TEAD•YAP/TAZ Transcription Factor Complex by Heterobifunctional Small Molecules that Bind to the TEAD Allosteric Lipid Pocket"

Department of Biochemistry

University of Illinois Urbana Champaign

600 S. Matthews Ave.

Urbana, IL 61801

### [1]H NMR (400 MHz) of compound TED-650

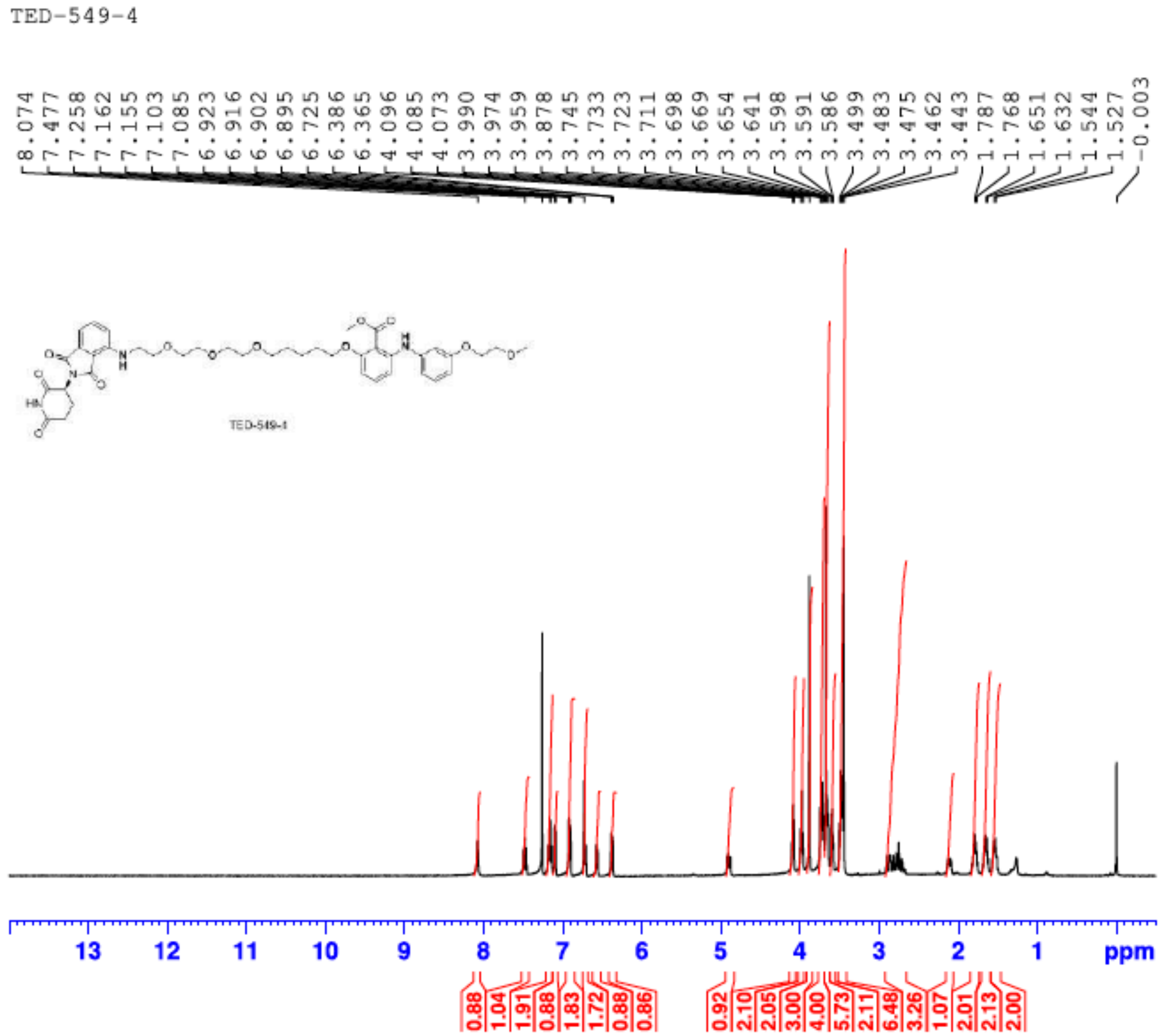

[1]H NMR (400 MHz) of compound TED-651

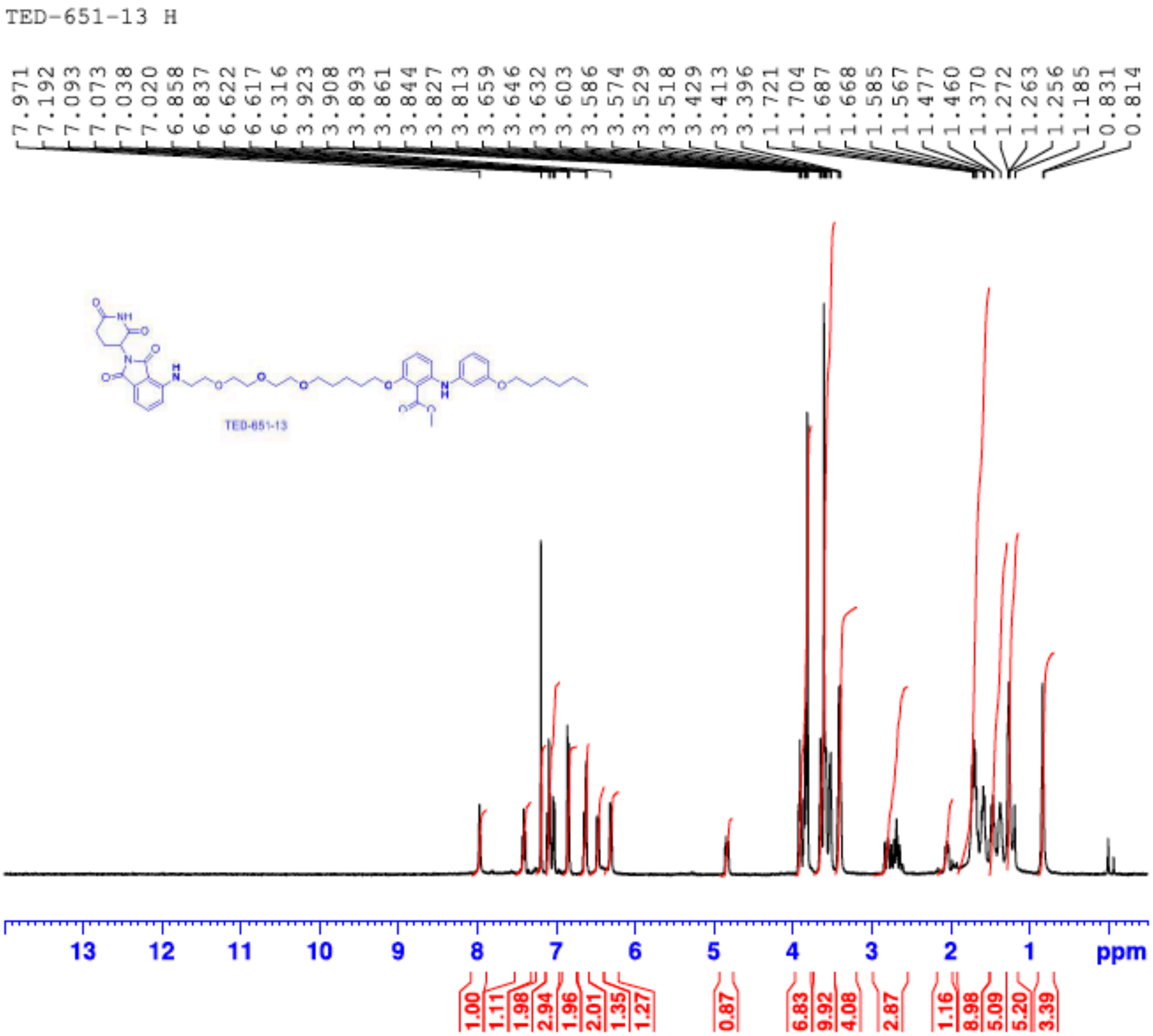

**$^{13}C$ NMR (100 MHz) of compound TED-651**

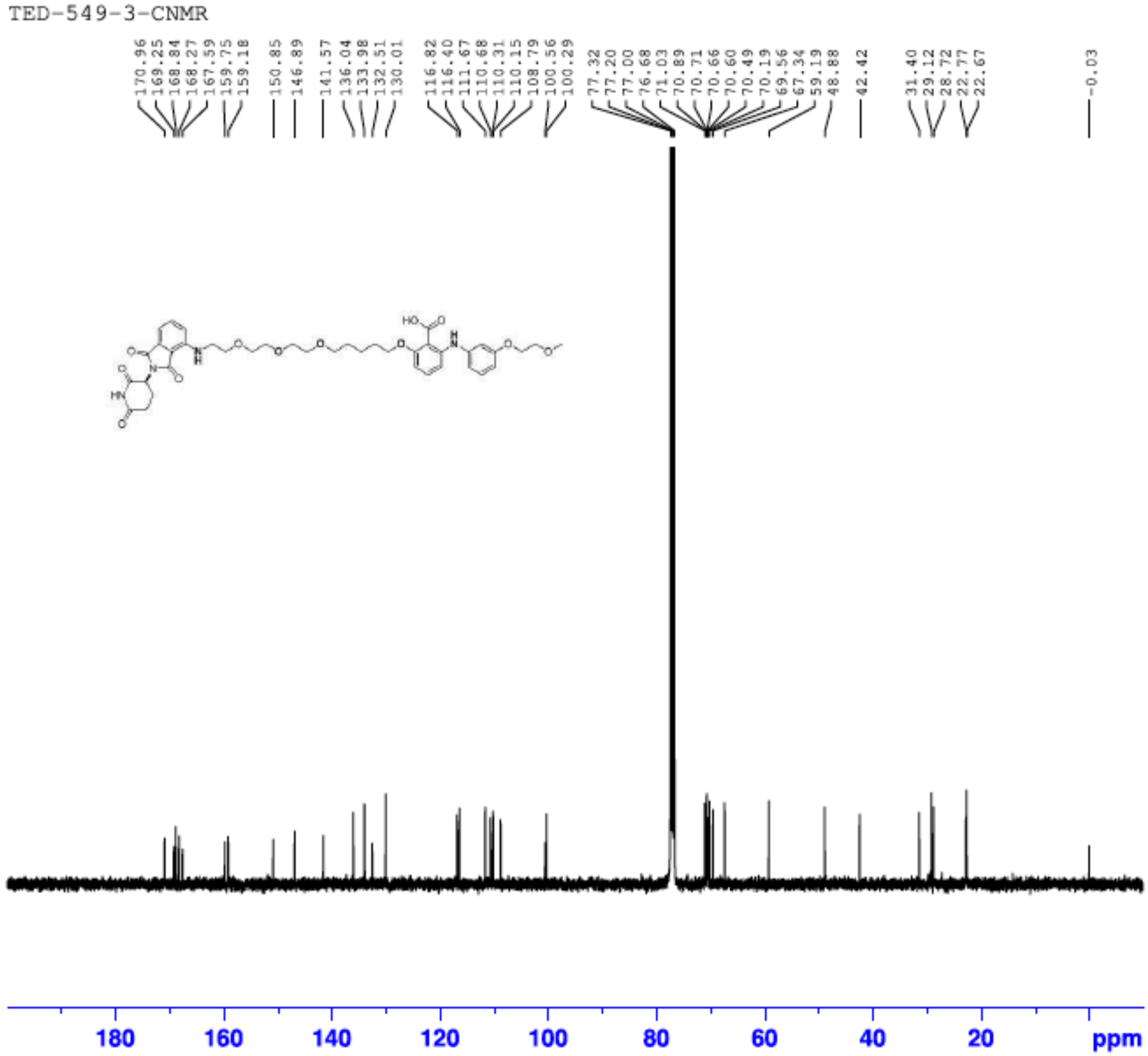

[1]H NMR (400 MHz) of compound TED-652

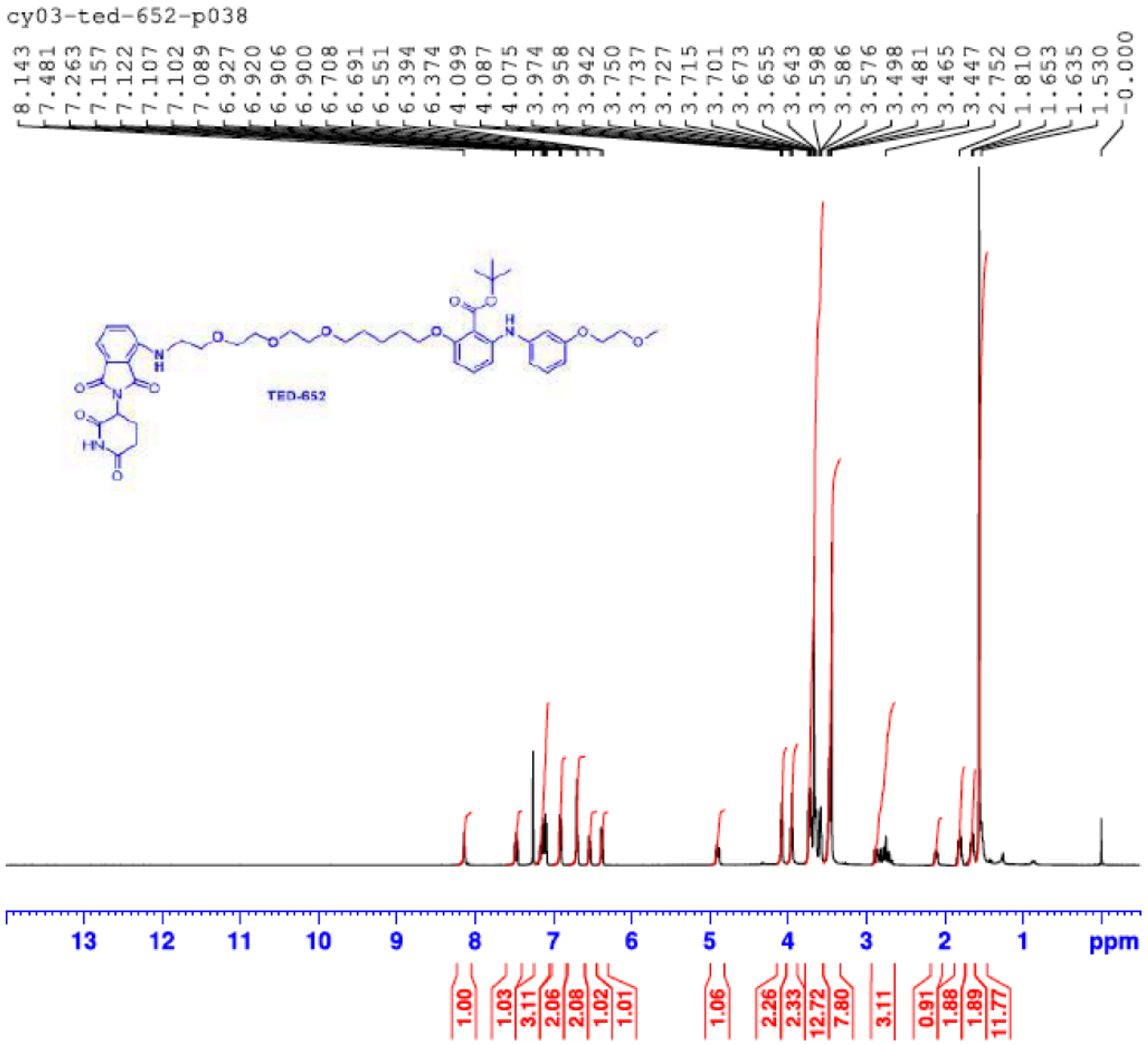

**$^{13}$C NMR (100 MHz) of compound TED-652**

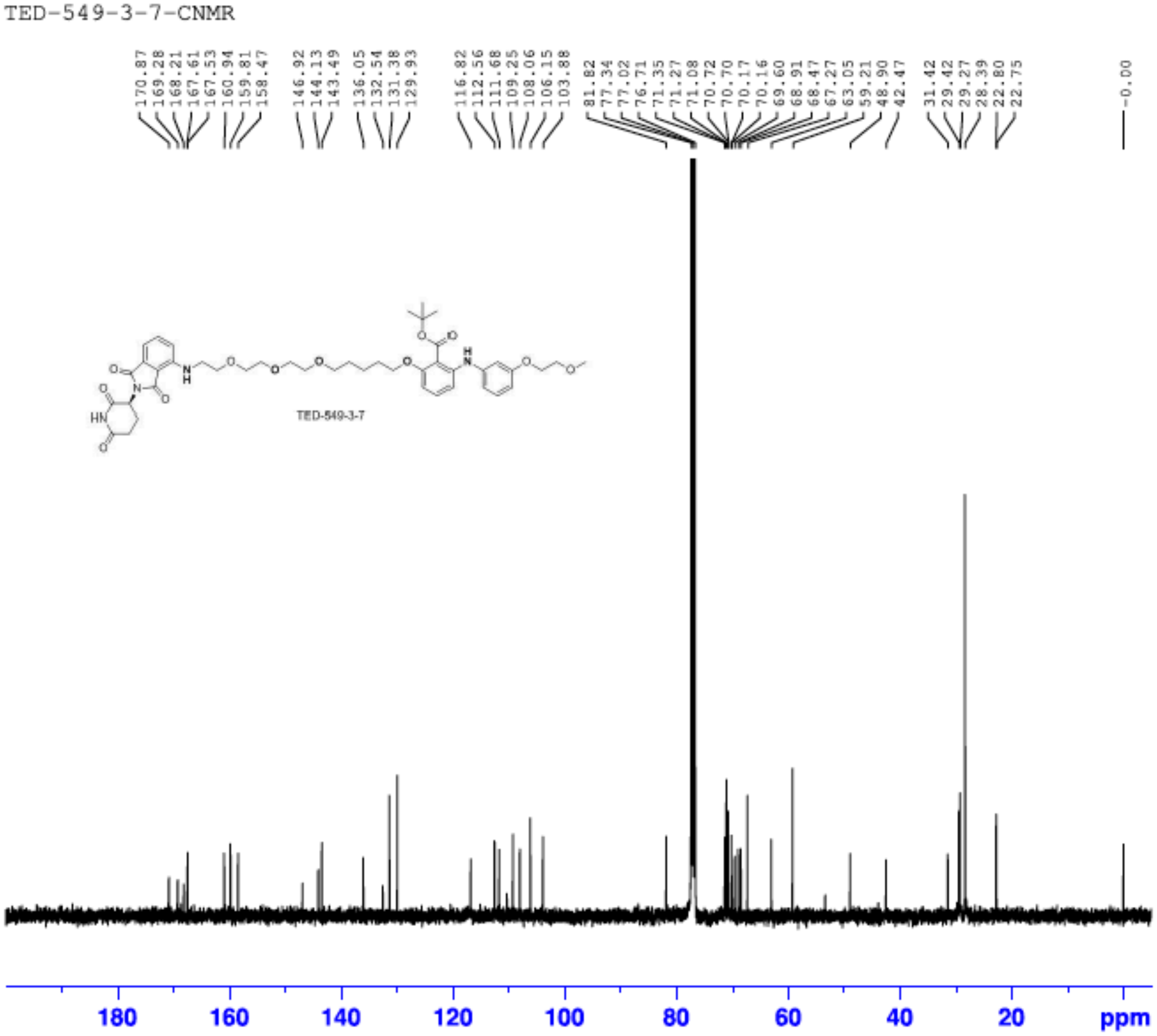

**[1]H NMR (400 MHz) of compound TED-670**

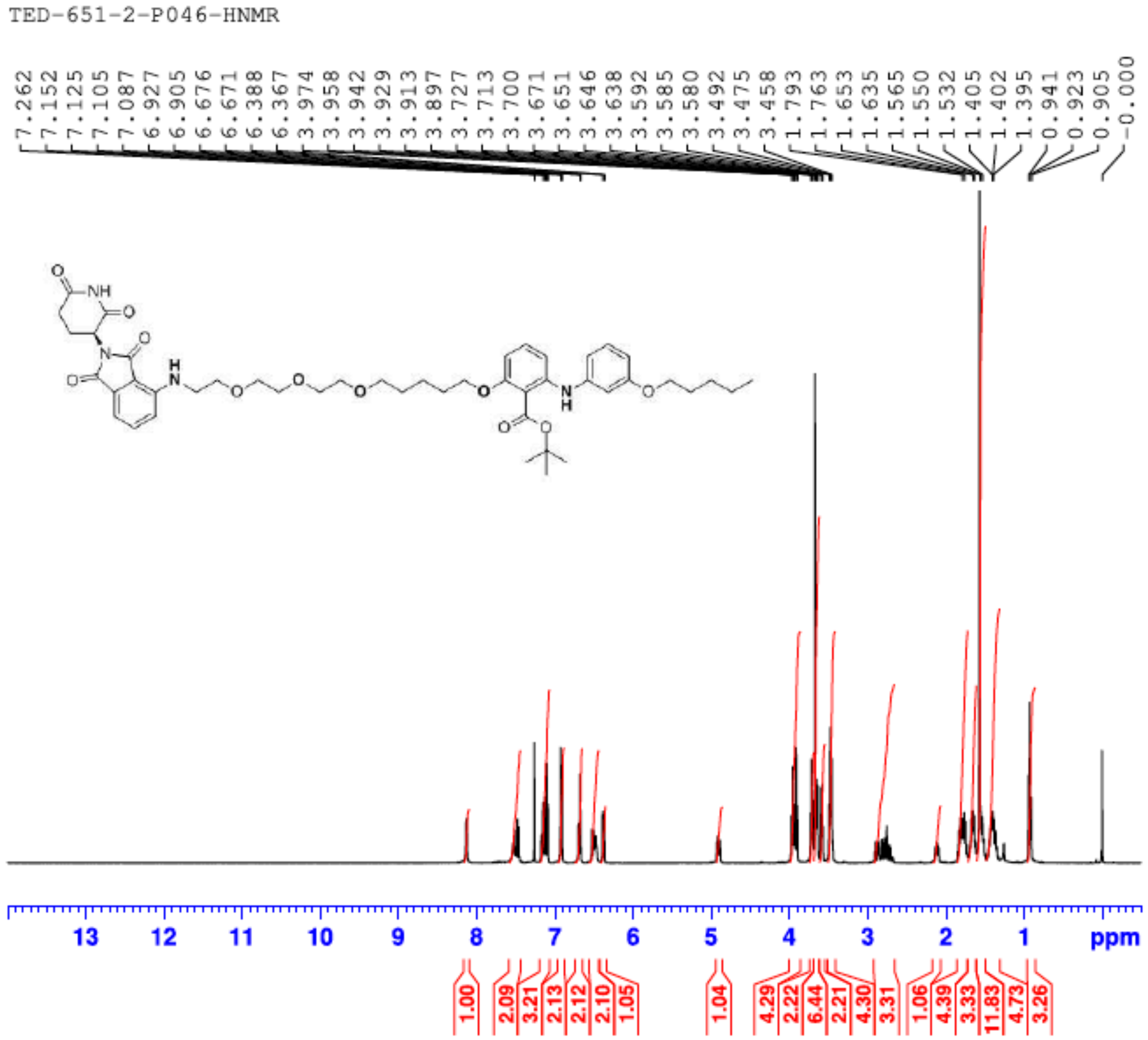

**$^{13}$C NMR (100 MHz) of compound TED-670**

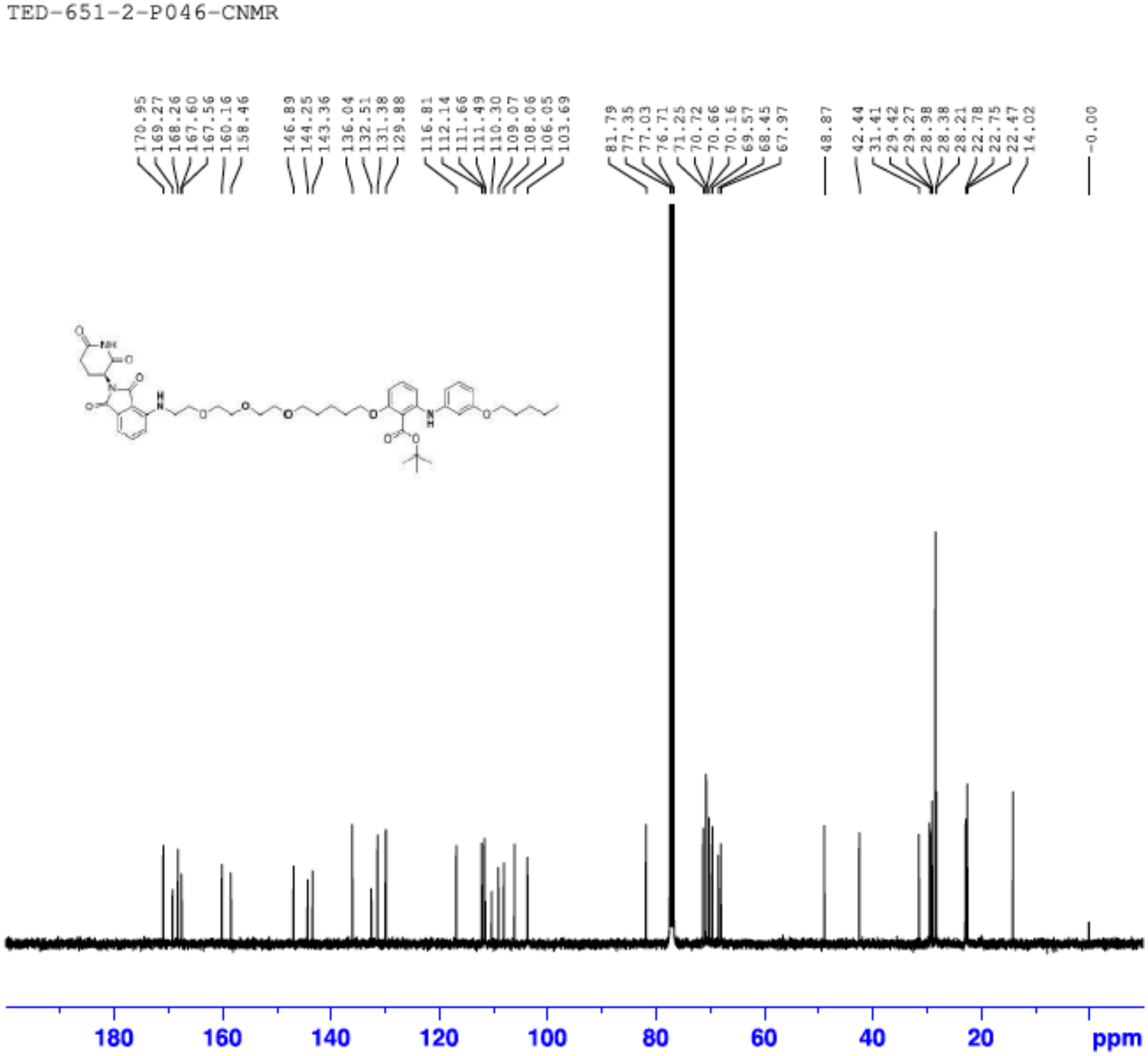

### [1]H NMR (400 MHz) of compound TED-671

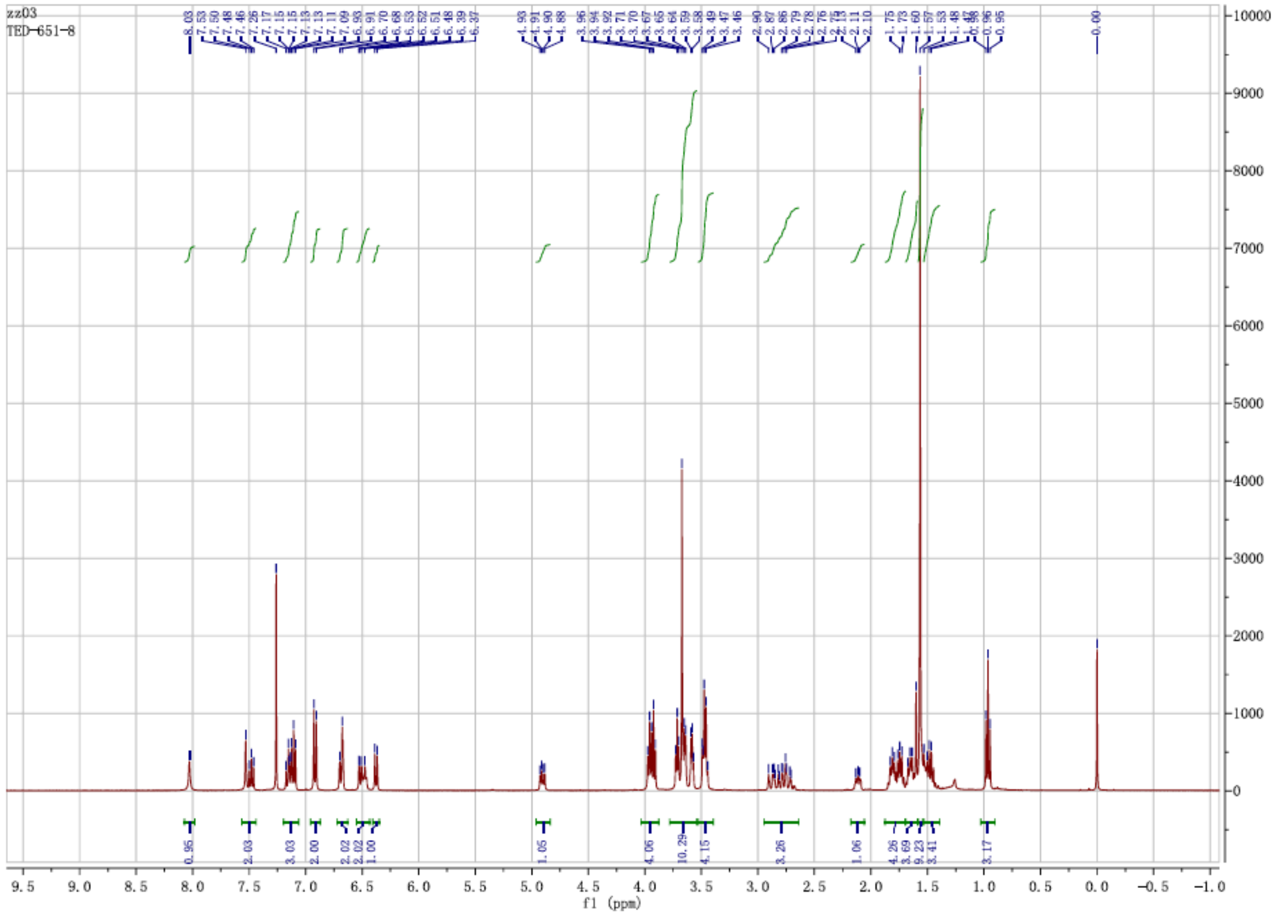

#### $^{13}$C NMR (100 MHz) of compound TED-671

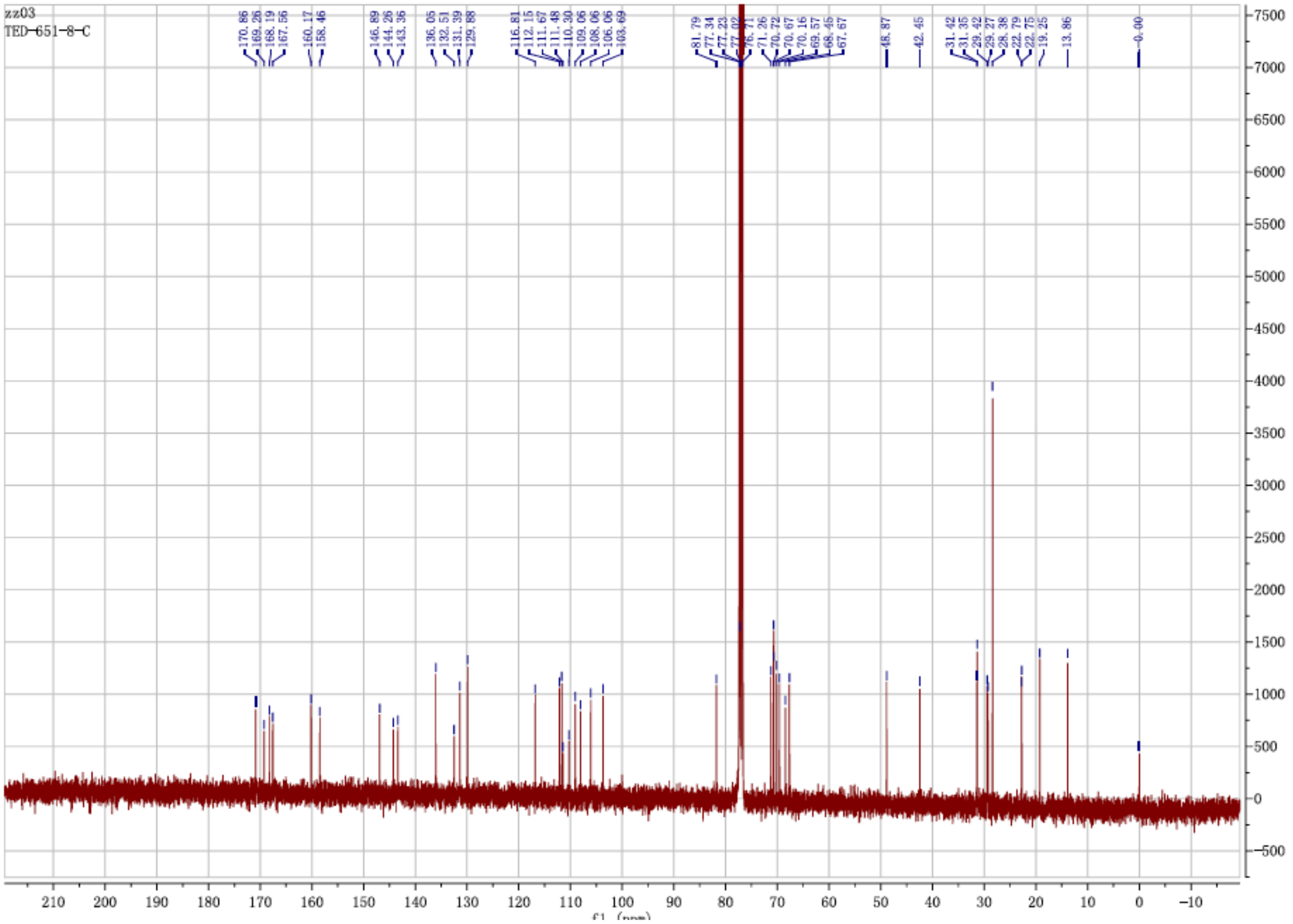

[1]H NMR (400 MHz) of compound TED-672

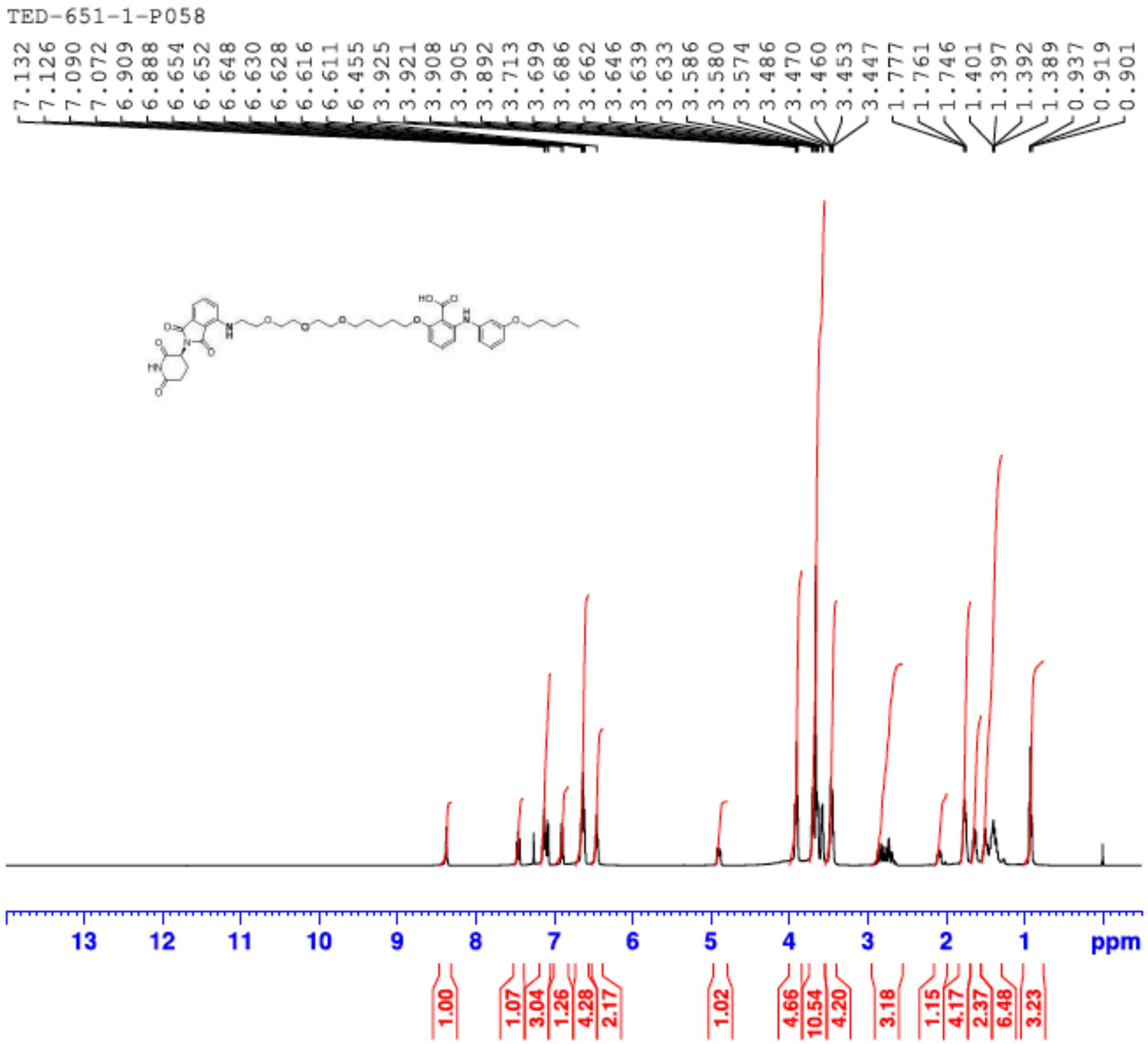

$^{13}$C NMR (100 MHz) of compound TED-672

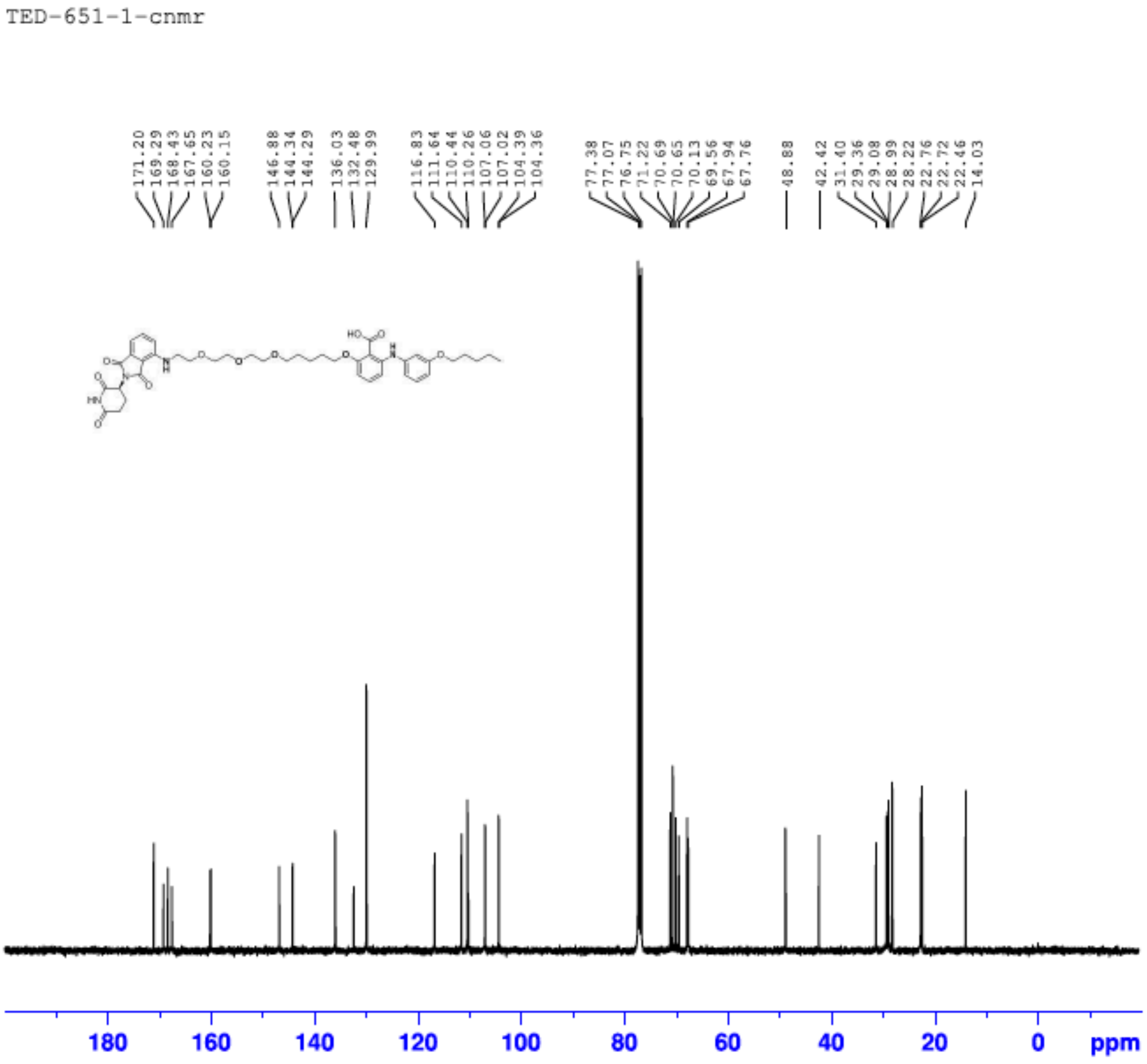

$^1$H NMR (400 MHz) of compound TED-673

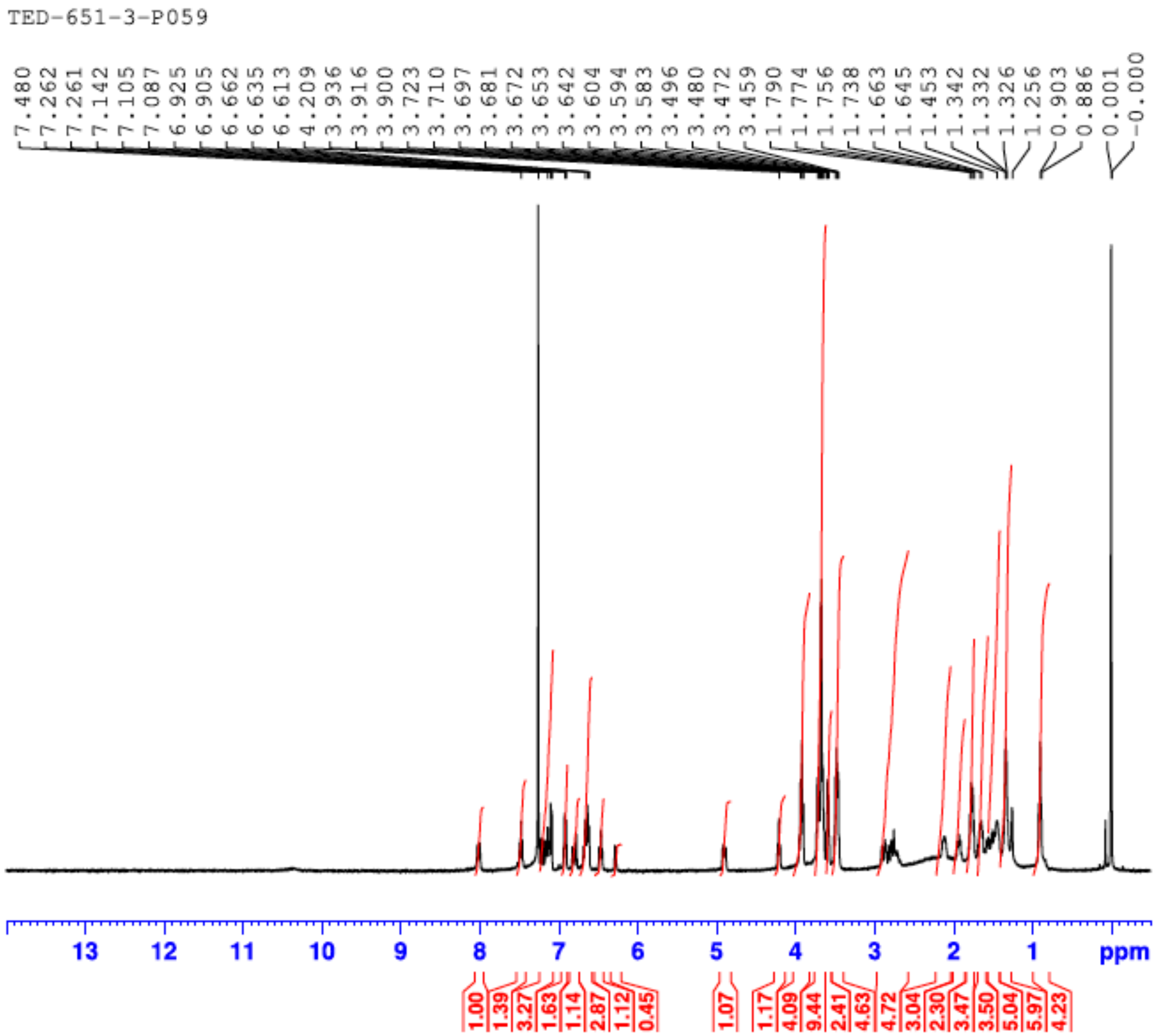

**$^1$H NMR (400 MHz) of compound TED-674**

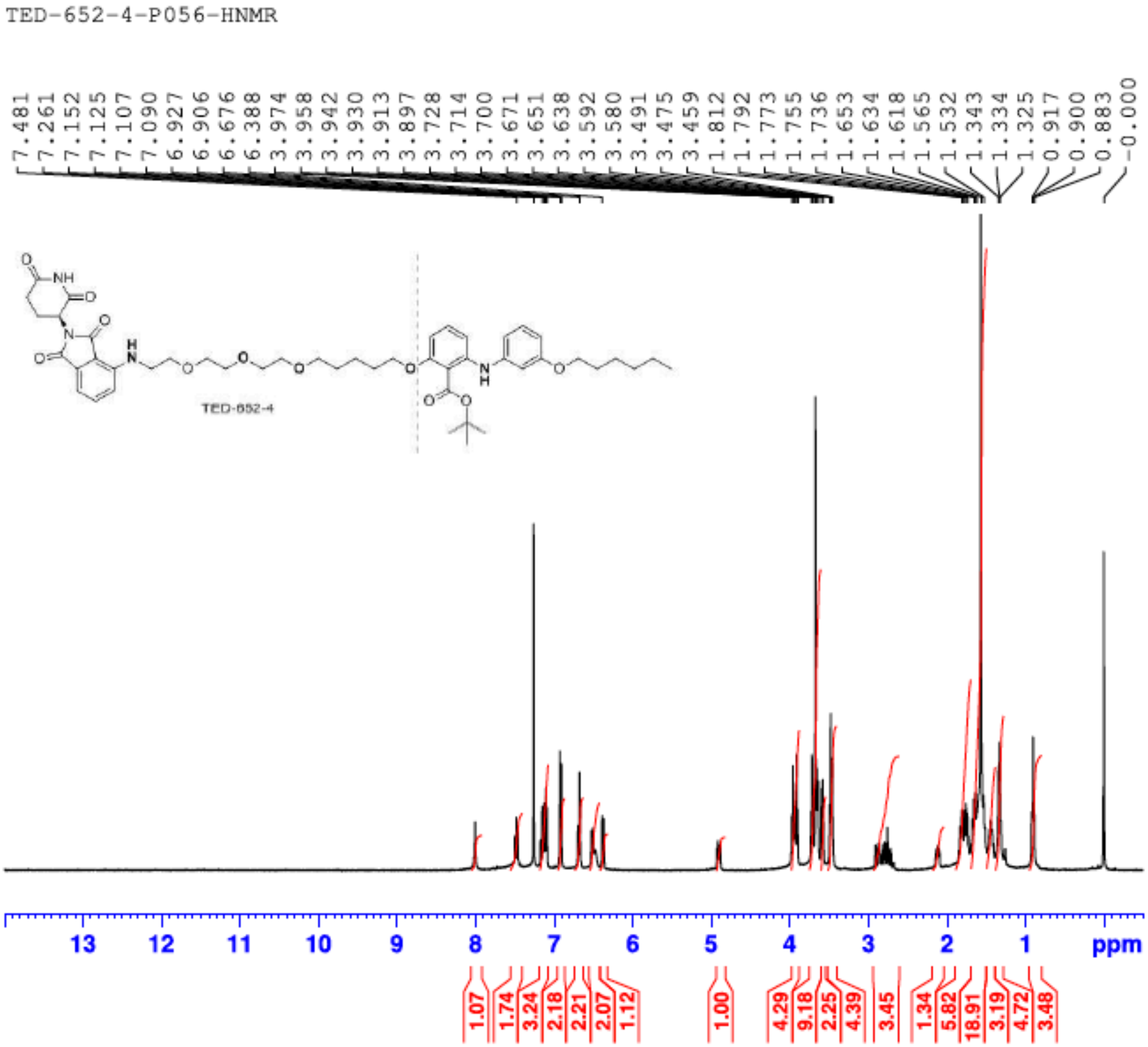

### $^{1}$H NMR (400 MHz) of compound TED-675

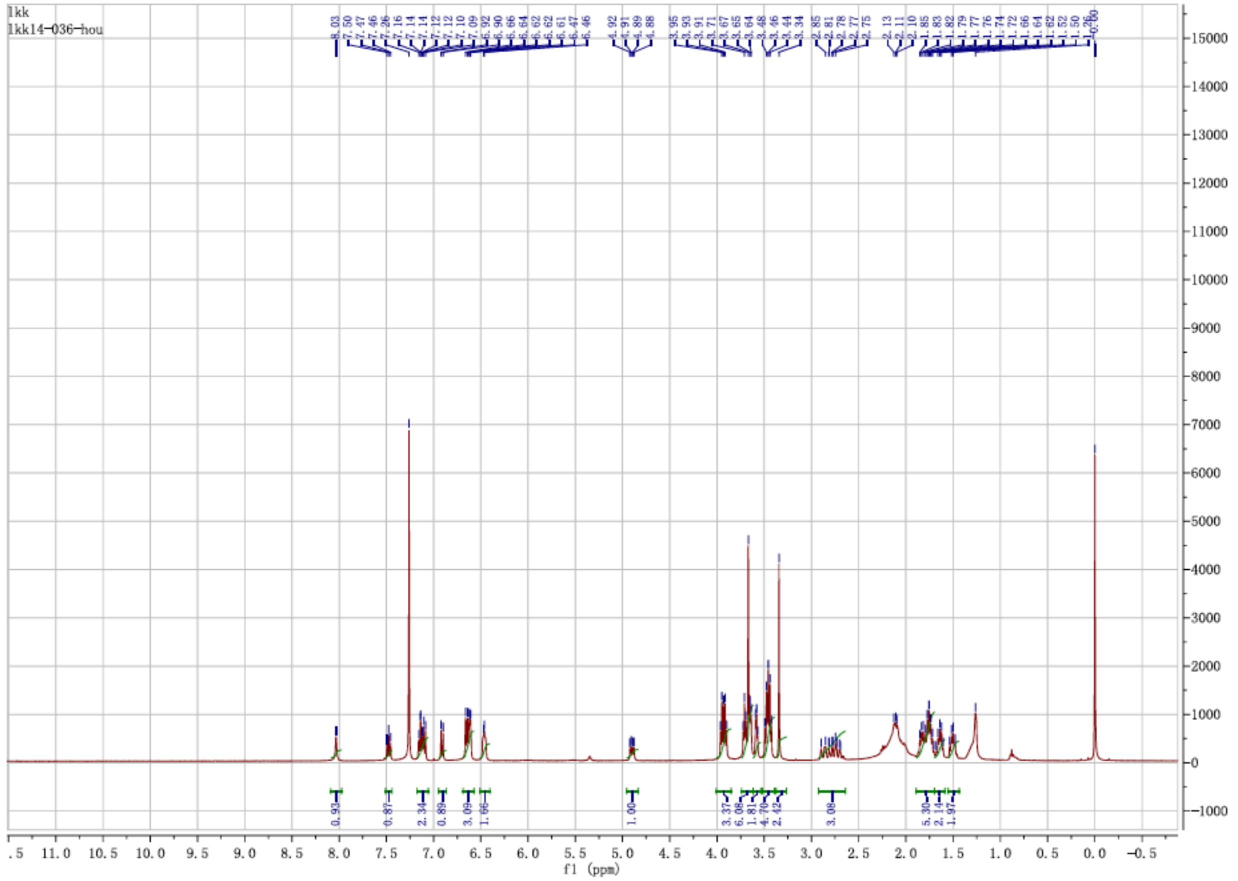

### $^1$H NMR (400 MHz) of compound TED-676

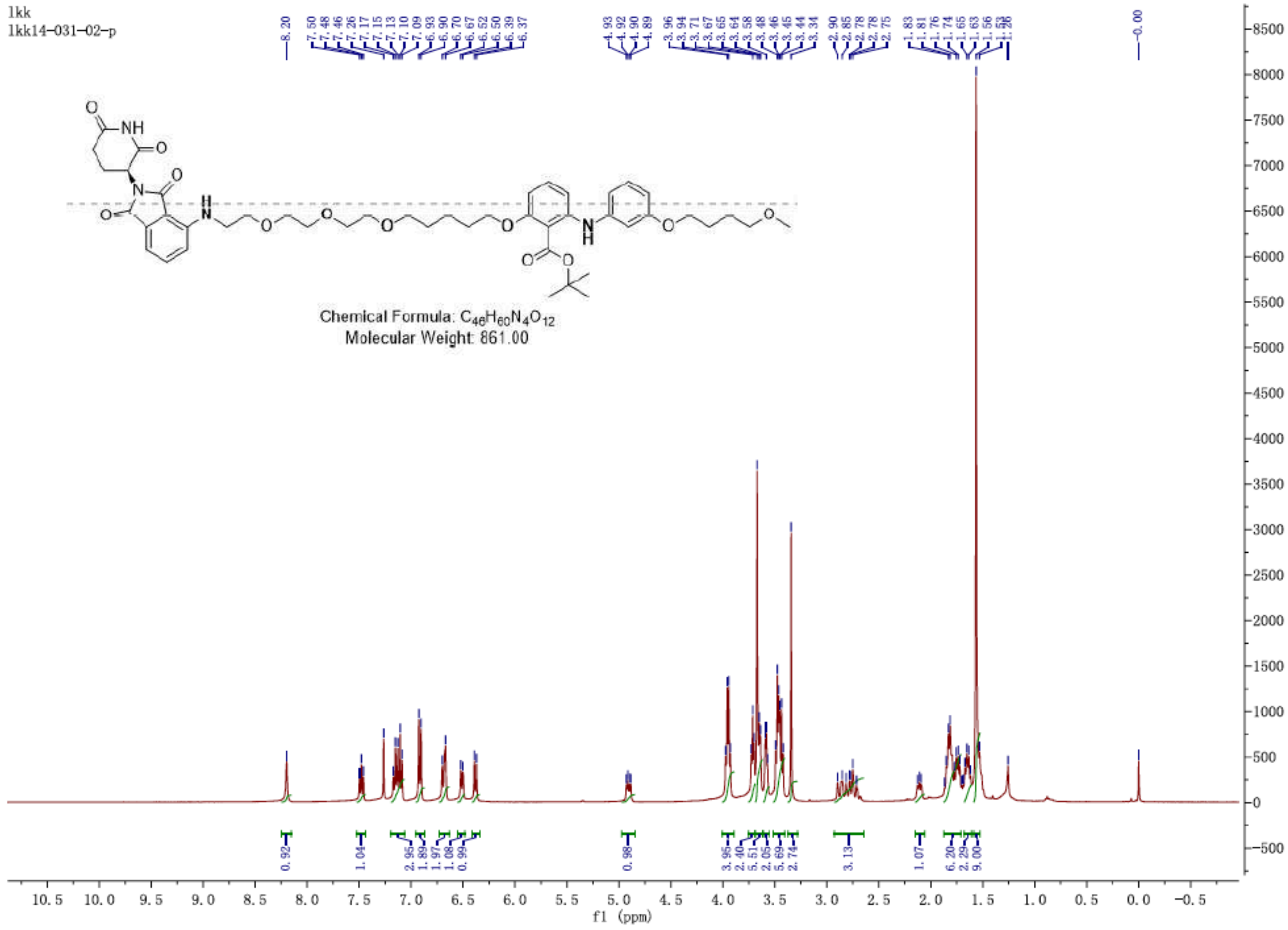

### $^1$H NMR (400 MHz) of compound TED-677

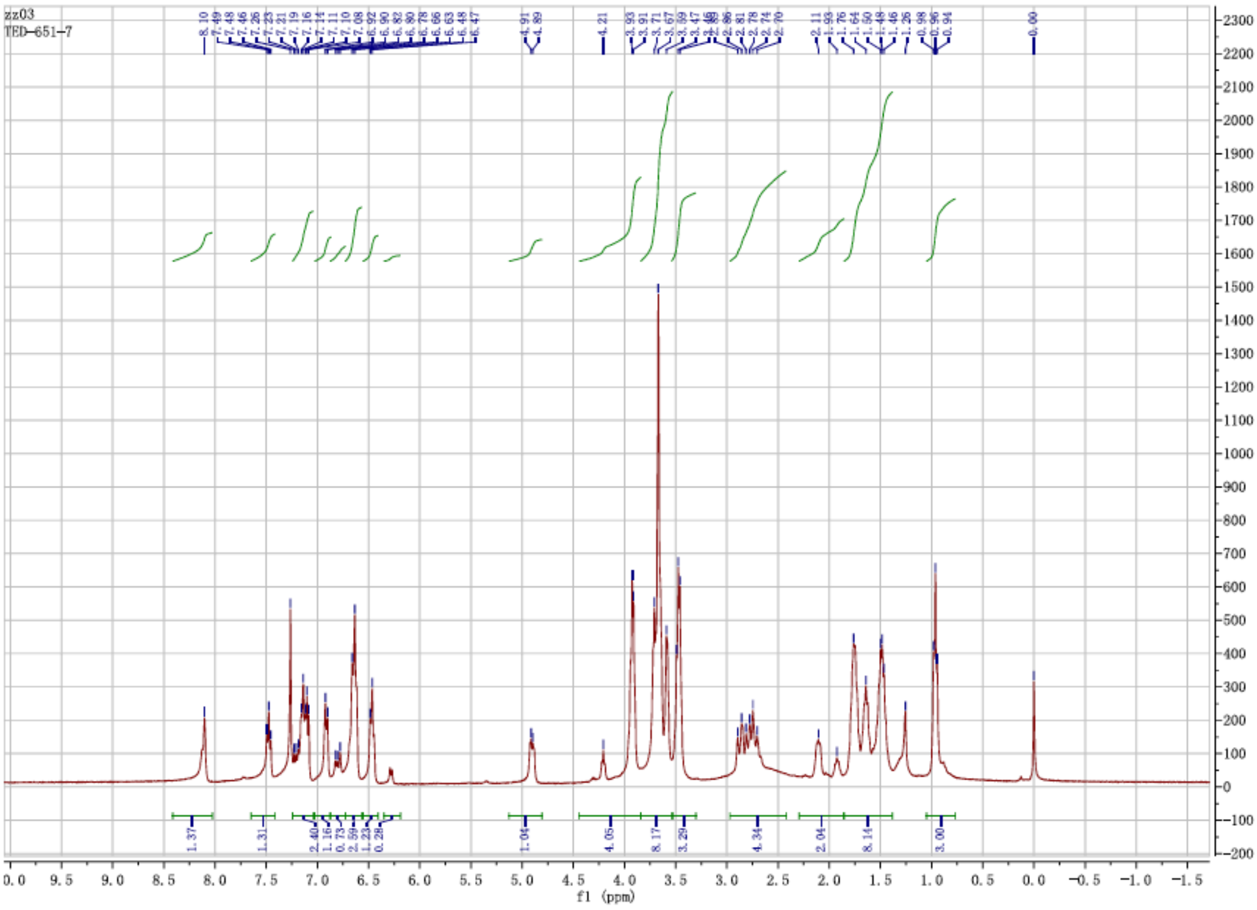

**[1]H NMR (400 MHz) of compound TED-688**

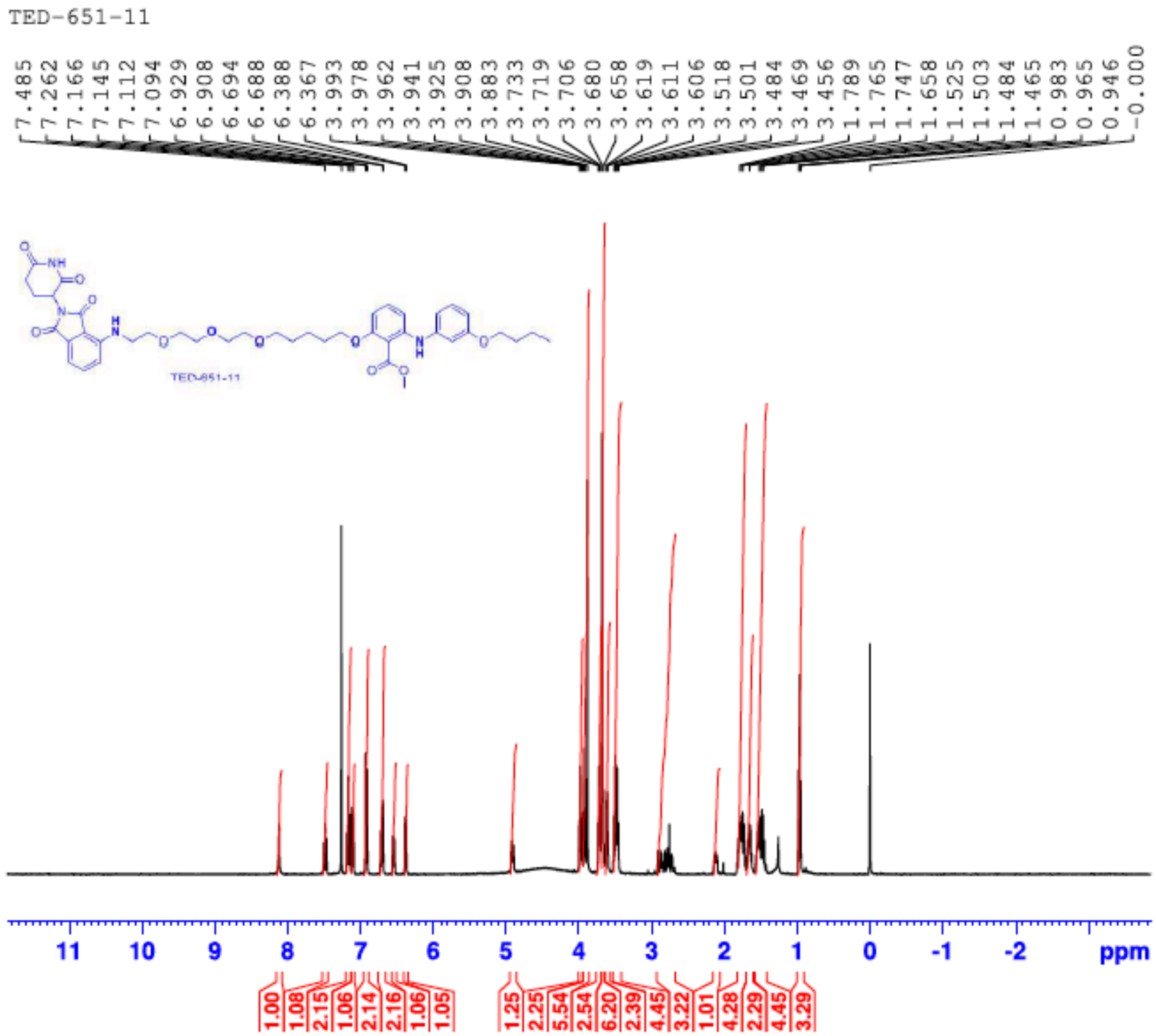

**[1]H NMR (400 MHz) of compound TED-689**

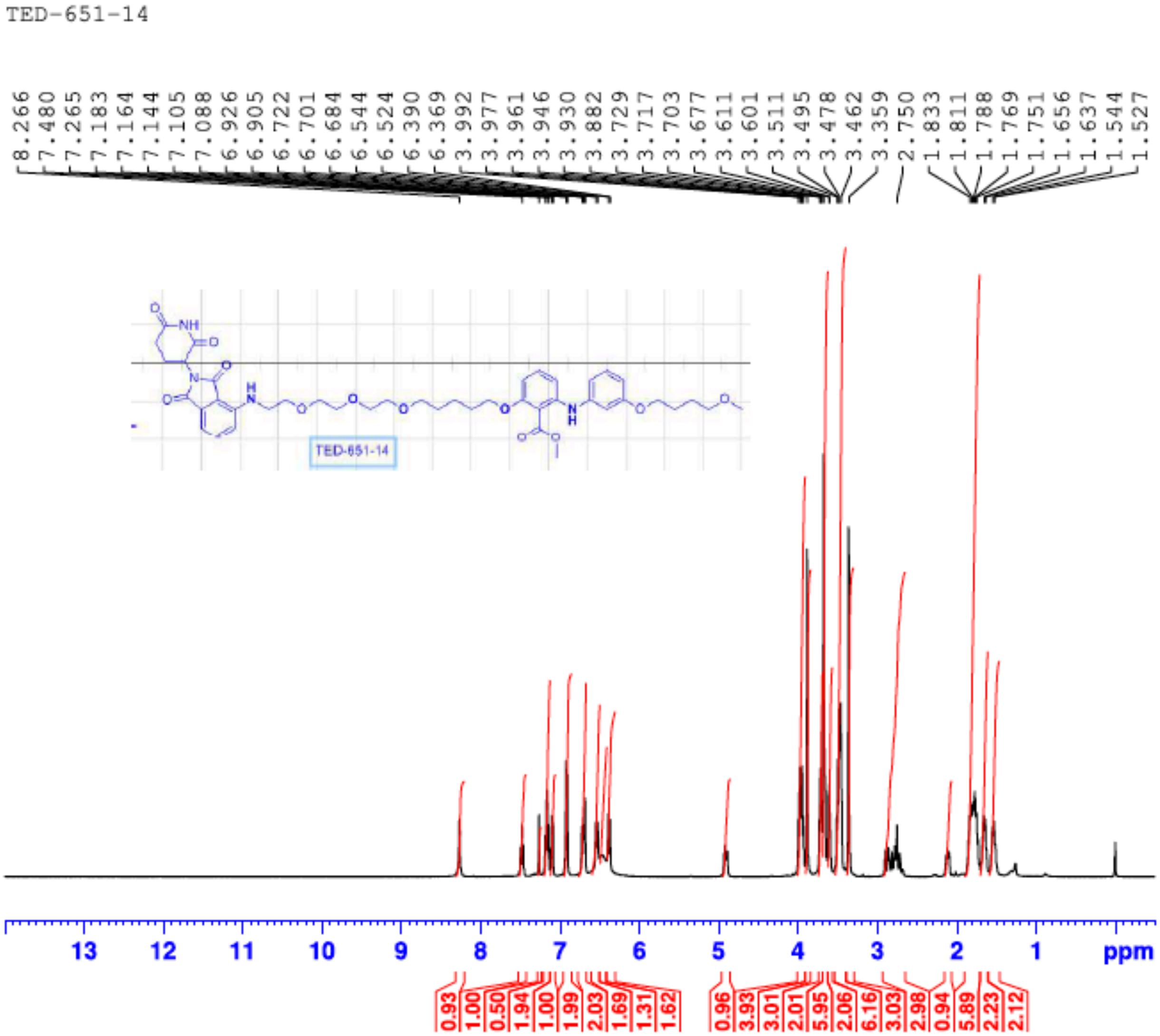

**[13]C NMR (100 MHz) of compound TED-689**

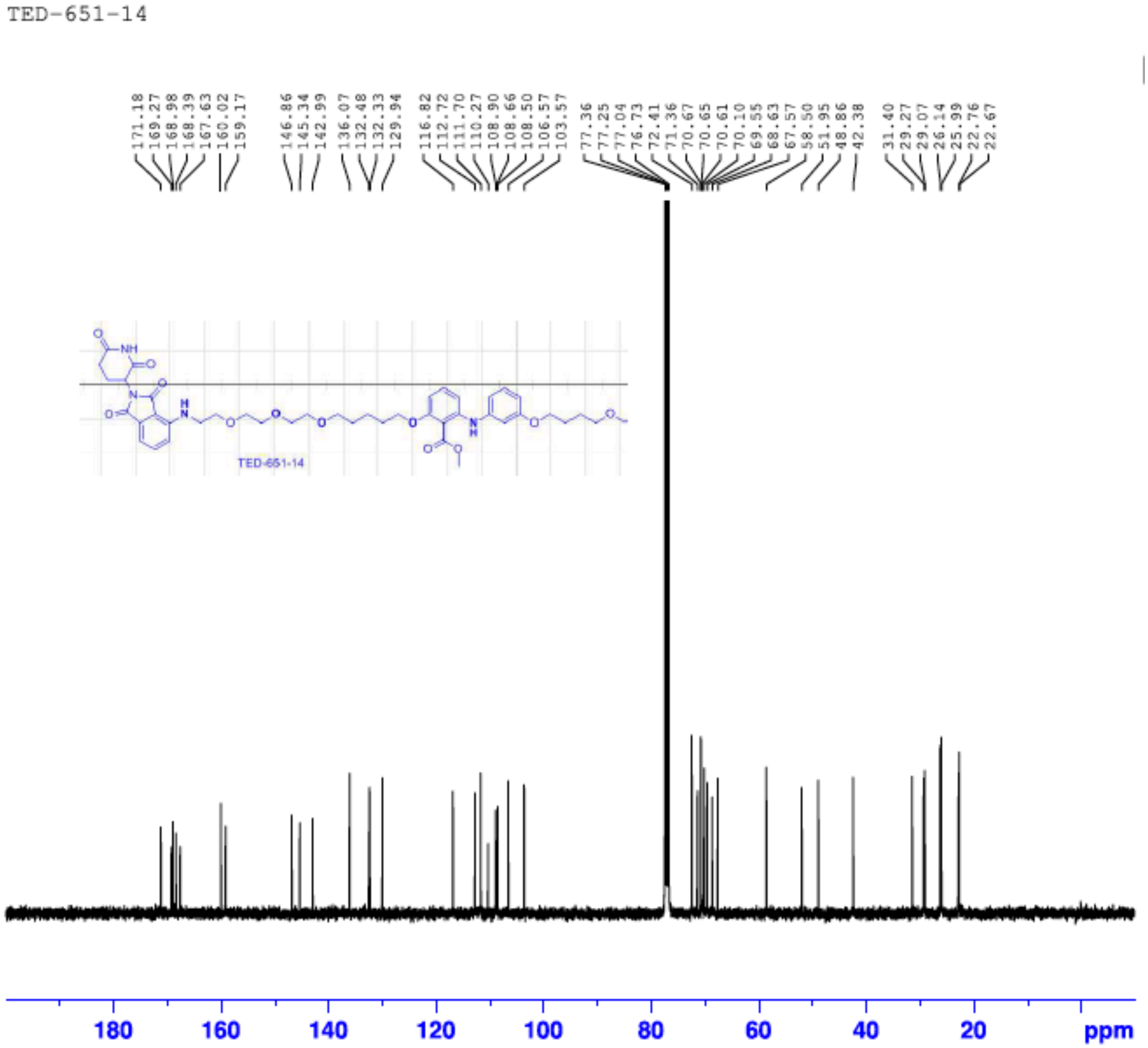

[1]H NMR (400 MHz) of compound TED-690

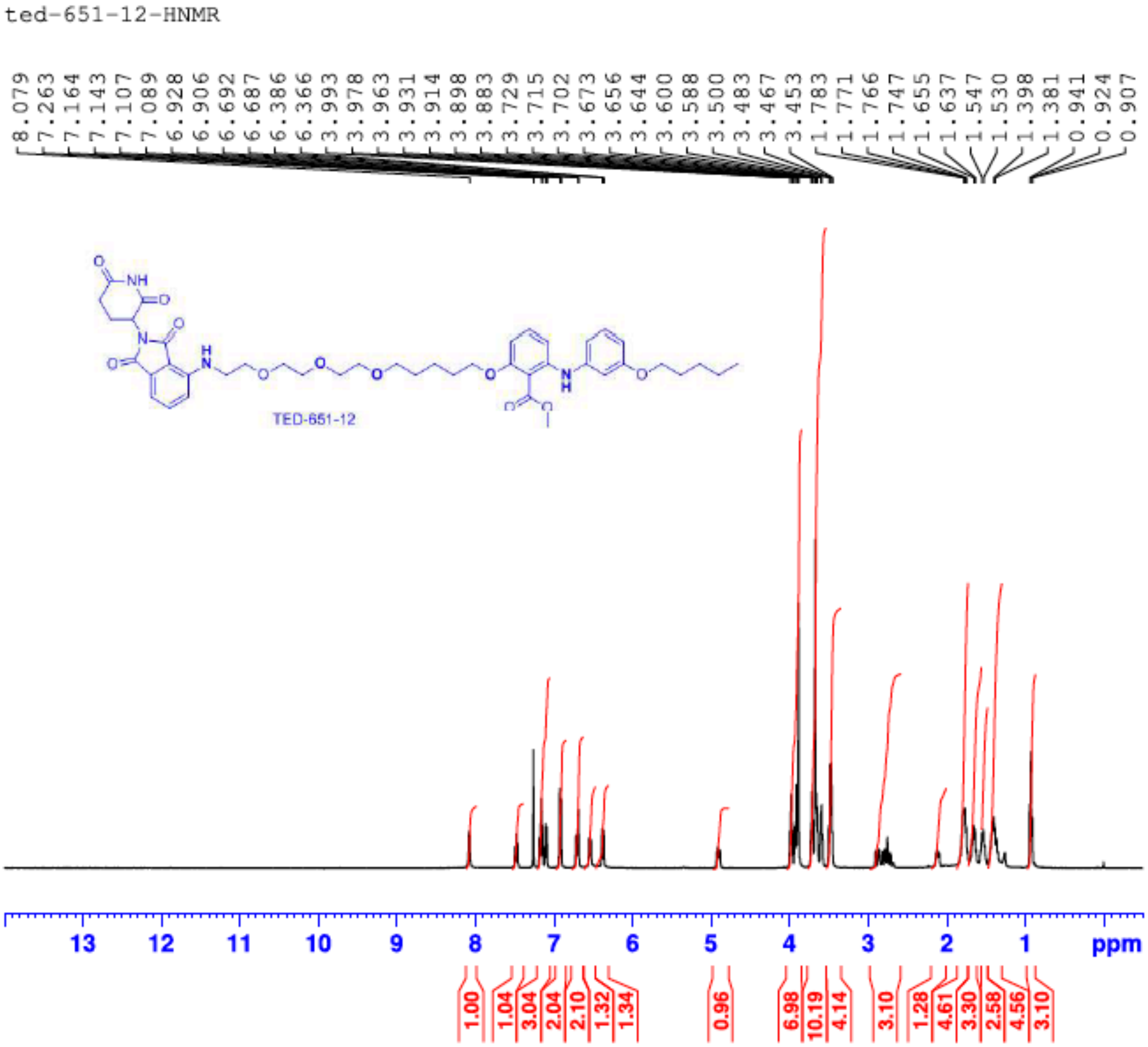

**[13]C NMR (100 MHz) of compound TED-690**

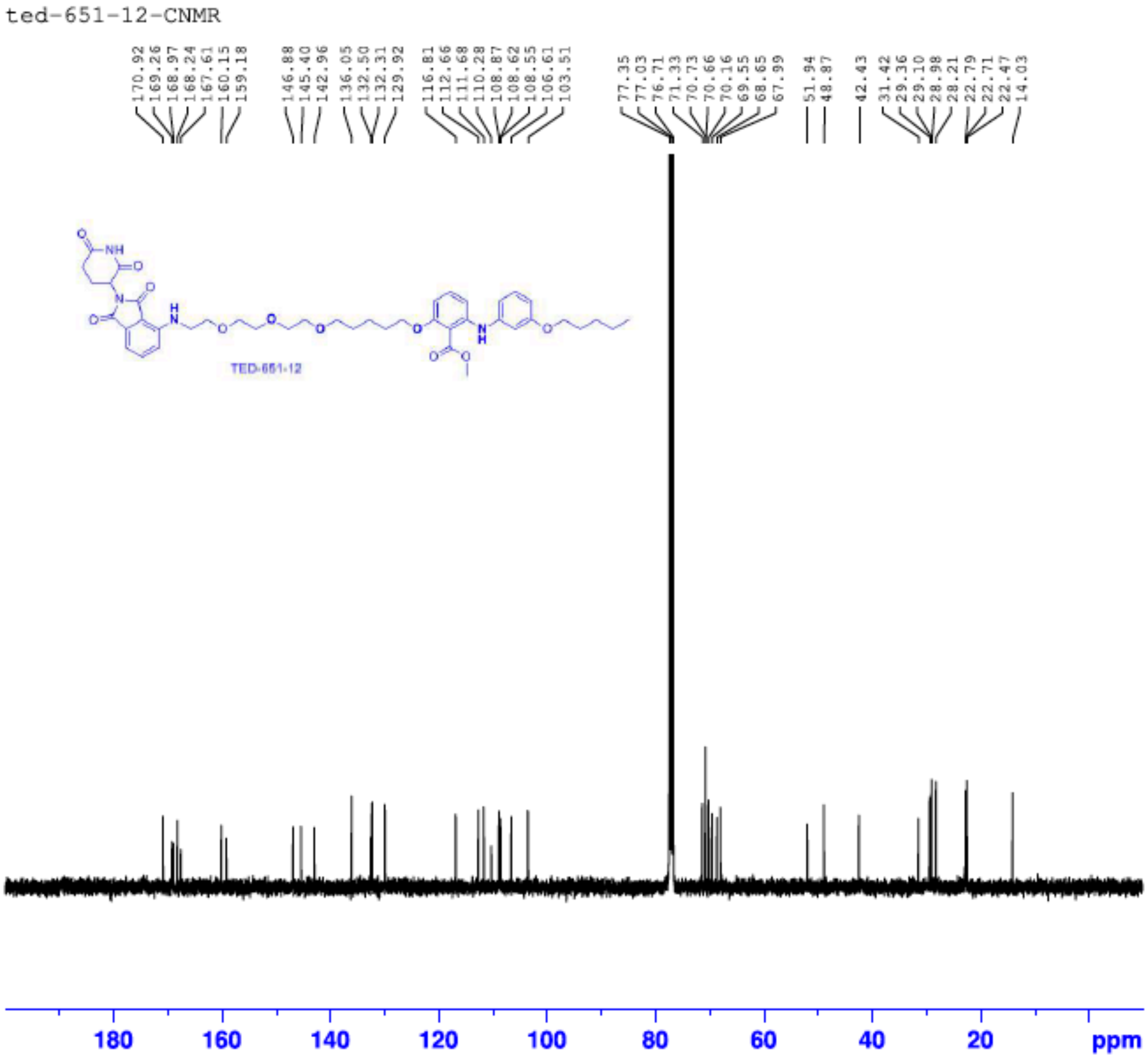

**$^{1}H$ NMR (400 MHz) of compound TED-734**

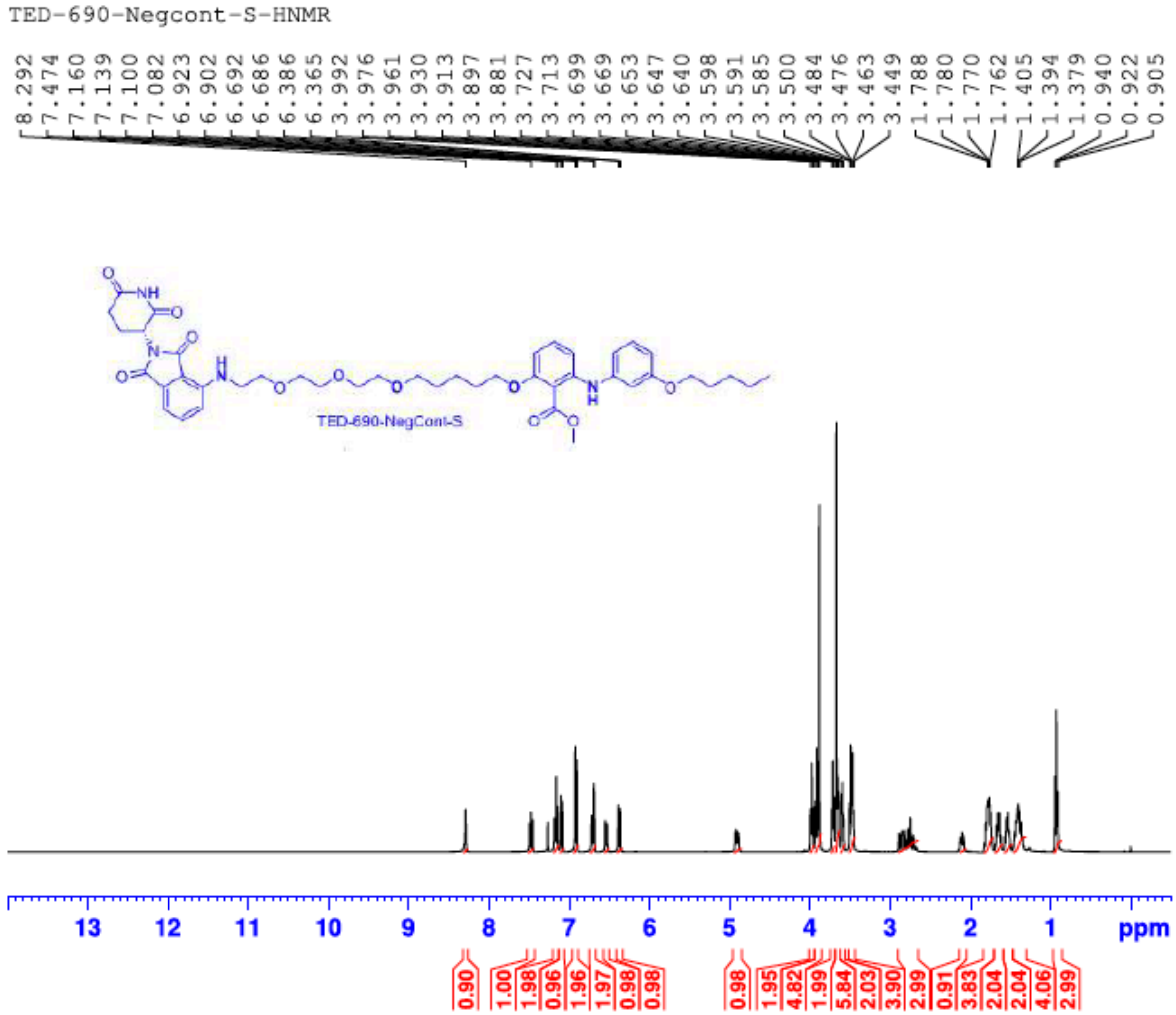

#### [13]C NMR (100 MHz) of compound TED-734

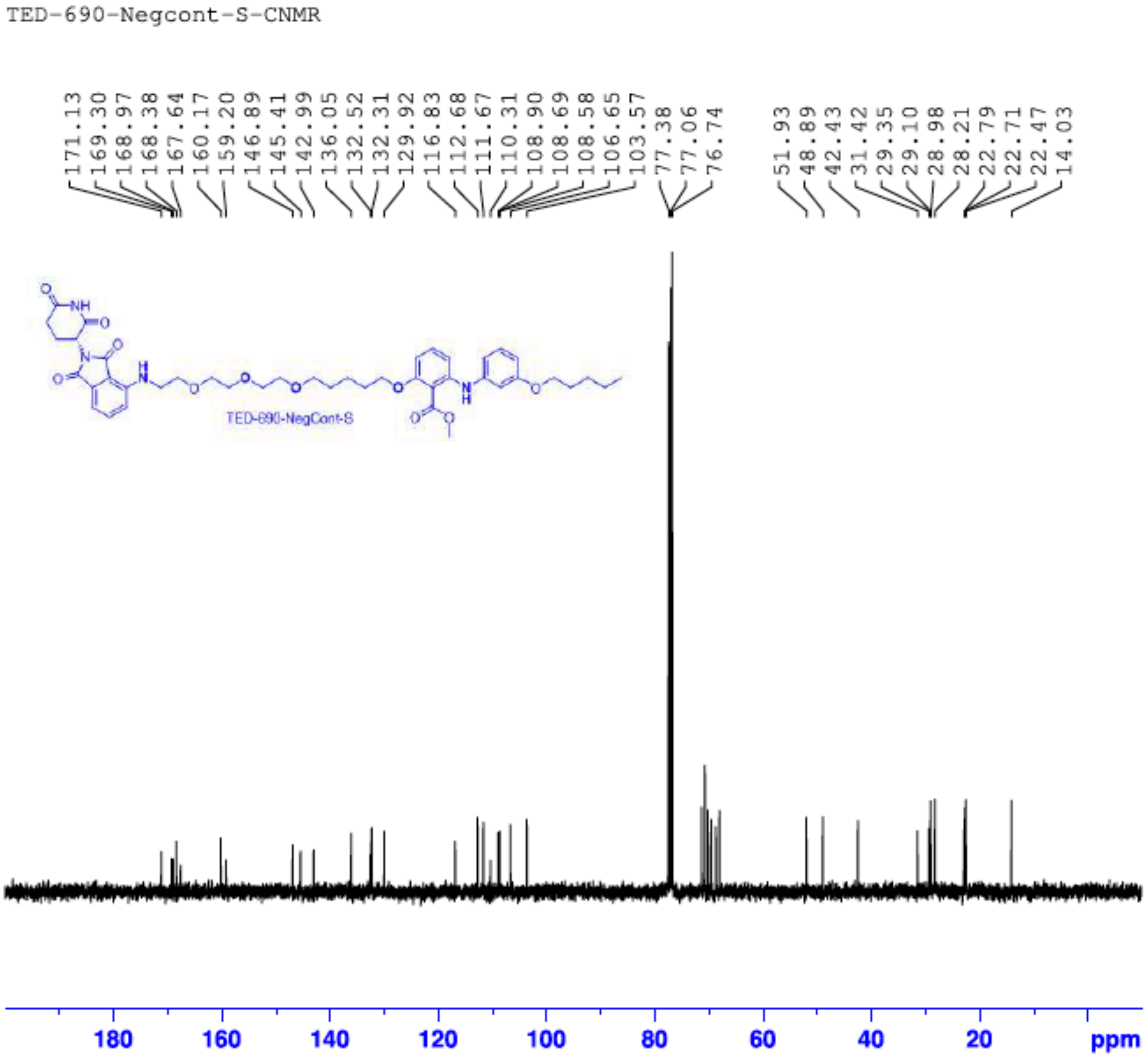

**HPLC Traces:**

**HPLC TED-650**

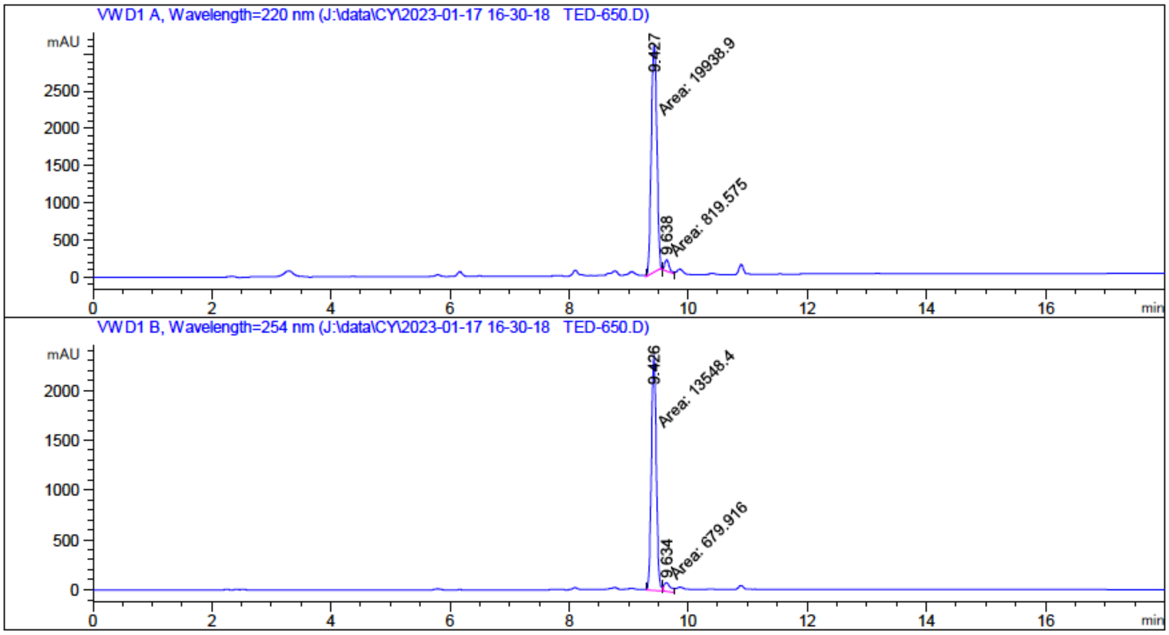


---

**UV 220 nm:**

**RT: 9.427 min** **Area %: 96.0518 %**

**RT: 9.638 min** **Area %: 3.9482 %**

**UV 254 nm:**

**RT: 9.426 min** **Area %: 95.2214 %**

**RT: 9.634 min** **Area %: 4.7786 %**

---

#### HPLC TED-651

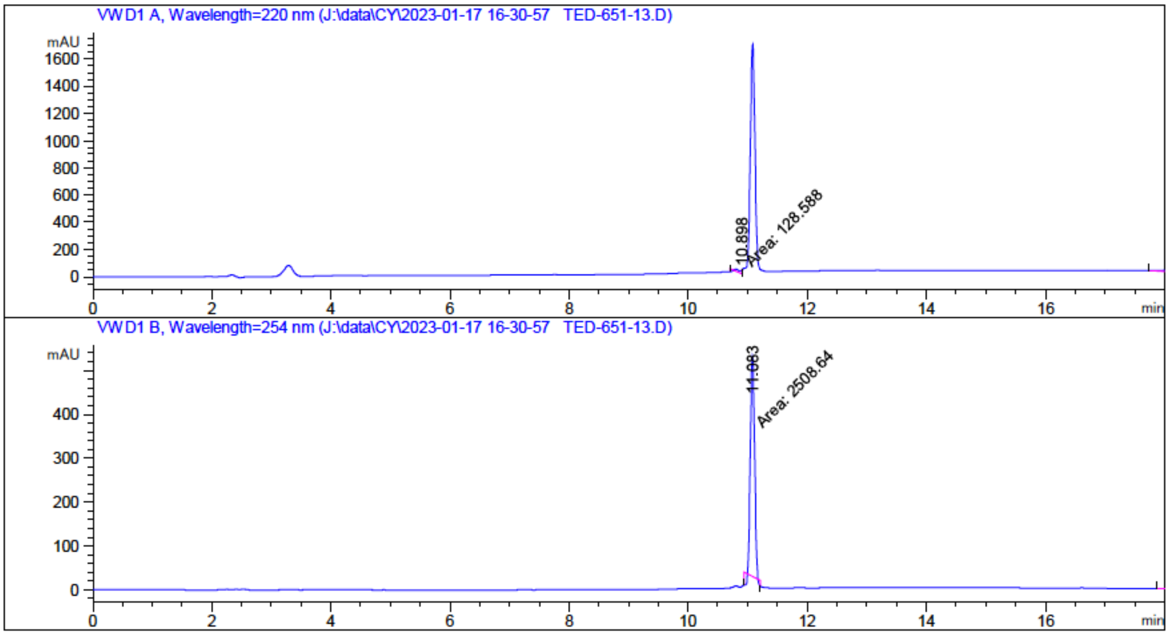


---

**UV 220 nm:**

**RT: 10.898 min** **Area %: 6.6483 %**

**RT: 18.130 min** **Area %: 93.3517 %**

**UV 254 nm:**

**RT: 11.083 min** **Area %: 96.1843 %**

**RT: 18.129 min** **Area %: 3.8157 %**

---

#### HPLC TED-652

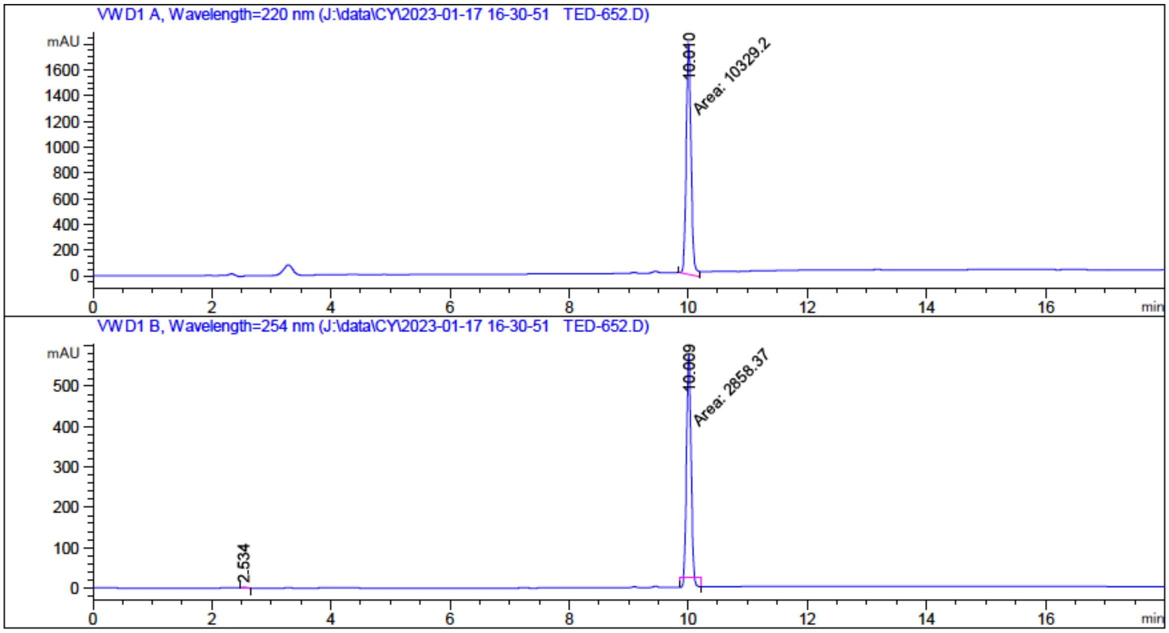


---

**UV 220 nm:**

**RT: 10.010 min** **Area %: 100.0000 %**

**UV 254 nm:**

**RT: 2.534 min** **Area %: 0.3371 %**

**RT: 10.009 min** **Area %: 99.6629 %**

---

**HPLC TED-670**

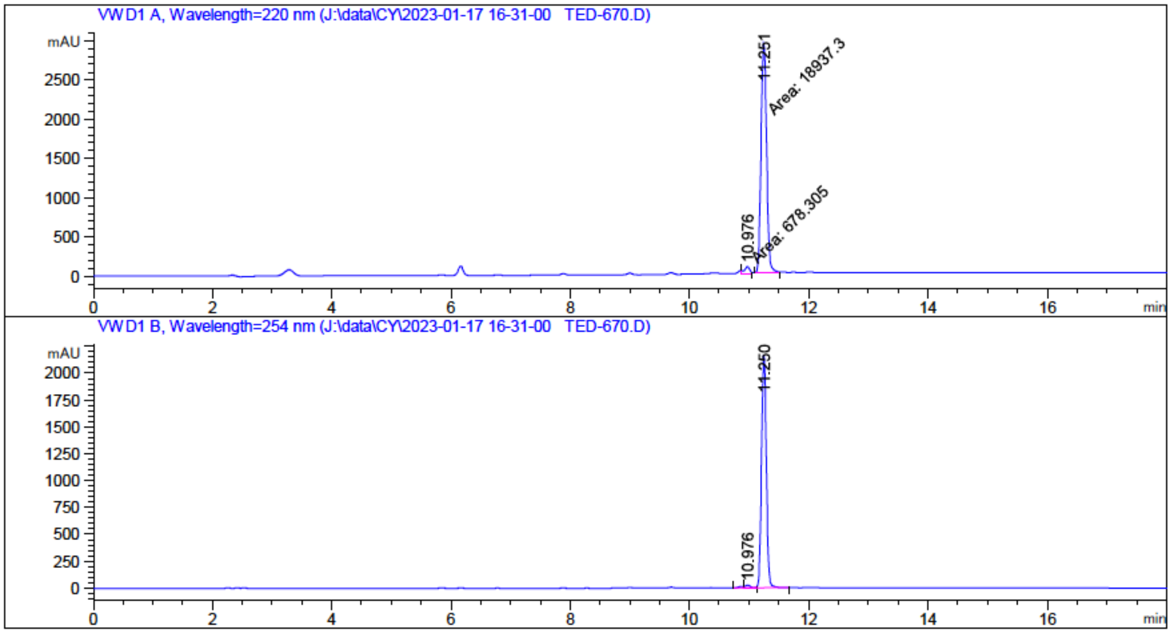


---

**UV 220 nm:**

**RT: 10.976 min** **Area %: 3.4580 %**

**RT: 11.251 min** **Area %: 96.5420 %**

**UV 254 nm:**

**RT: 10.976 min** **Area %: 1.2236 %**

**RT: 11.250 min** **Area %: 98.7764 %**

---

**HPLC TED-671**

---

**UV 220 nm:**

**RT: min Area %: %**

**UV 254 nm:**

**RT: min Area %: %**

---

**HPLC TED-672**

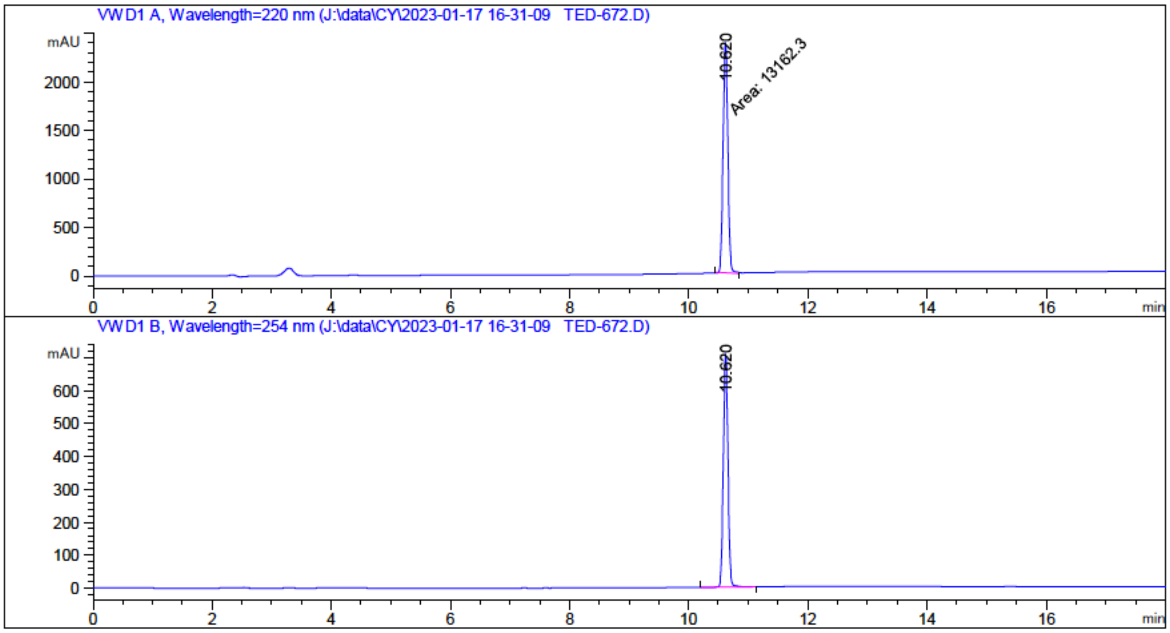


---

**UV 220 nm:**

**RT: 10.620 min Area %: 100.0000 %**

**UV 254 nm:**

**RT: 10.620 min Area %: 100.0000 %**

---

**HPLC TED-673**

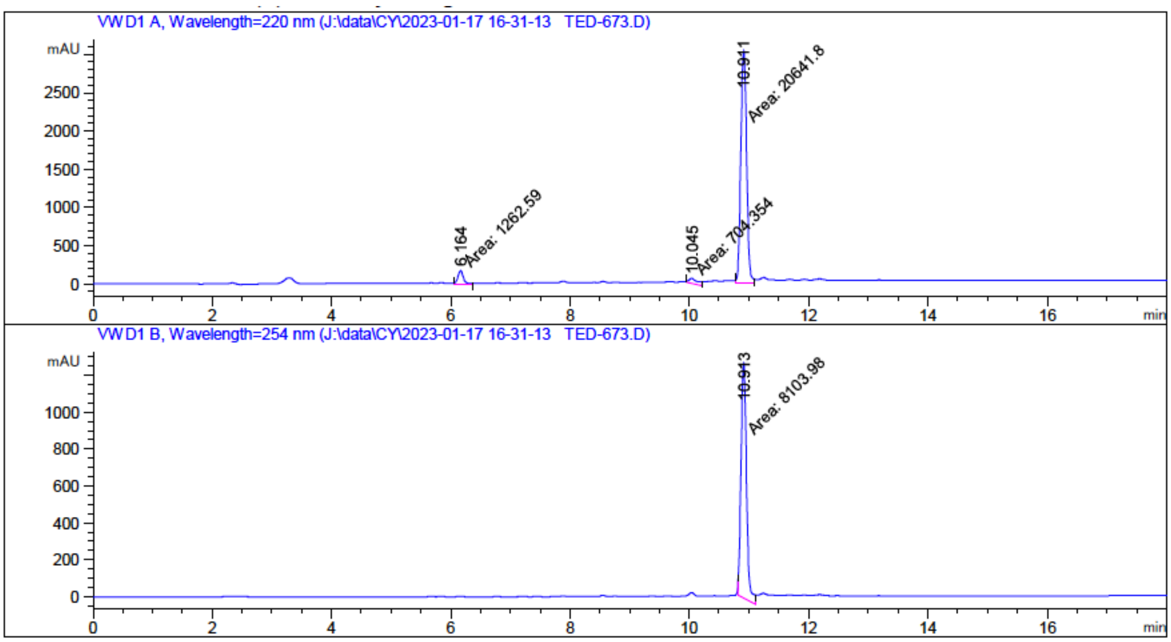


---

**UV 220 nm:**

**RT: 6.164 min** **Area %: 5.5845 %**

**RT: 10.045 min** **Area %: 3.1154 %**

**RT: 10.911 min** **Area %: 91.3001 %**

**UV 254 nm:**

**RT: 10.913 min** **Area %: 100.0000 %**

---

**HPLC TED-674**

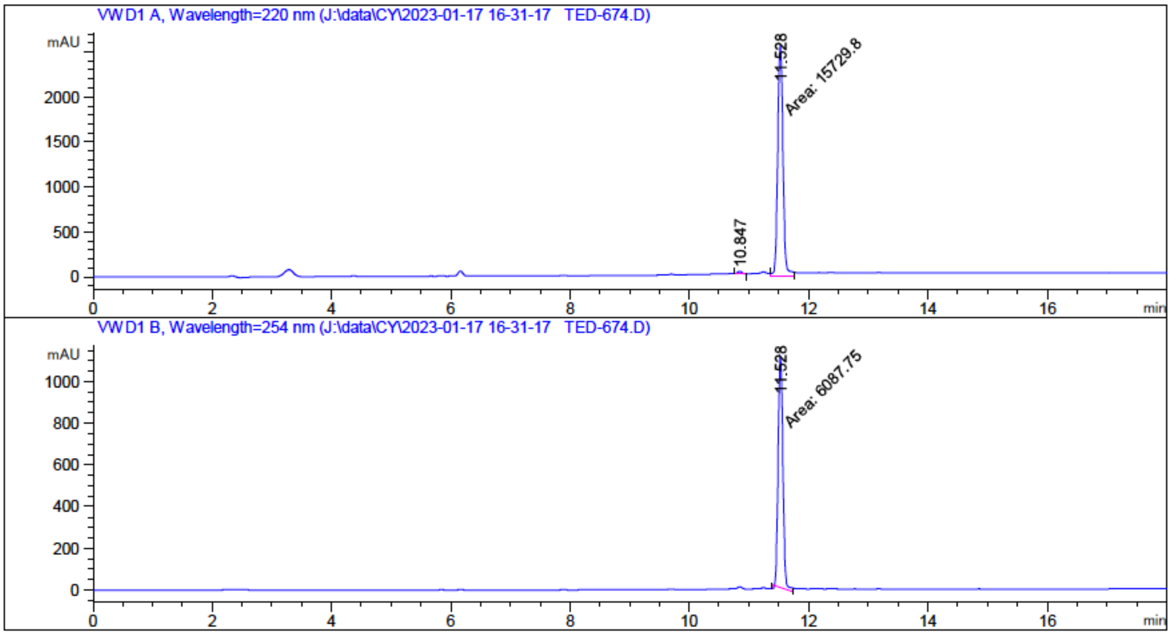


---

**UV 220 nm:**

**RT: 10.847 min** **Area %: 0.9279 %**

**RT: 11.528 min** **Area %: 99.0721 %**

**UV 254 nm:**

**RT: 11.528 min** **Area %: 100.0000 %**

---

**HPLC TED-688**

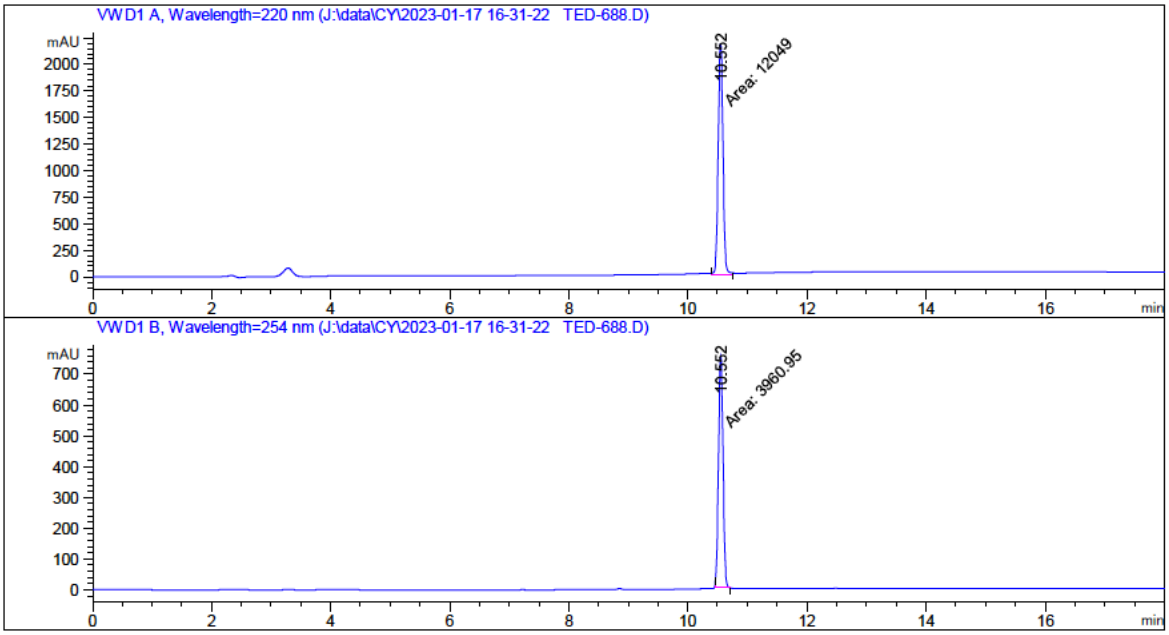


---

**UV 220 nm:**

**RT: 10.552 min** **Area %: 100.0000 %**

**UV 254 nm:**

**RT: 10.552 min** **Area %: 100.0000 %**

---

**HPLC TED-689**

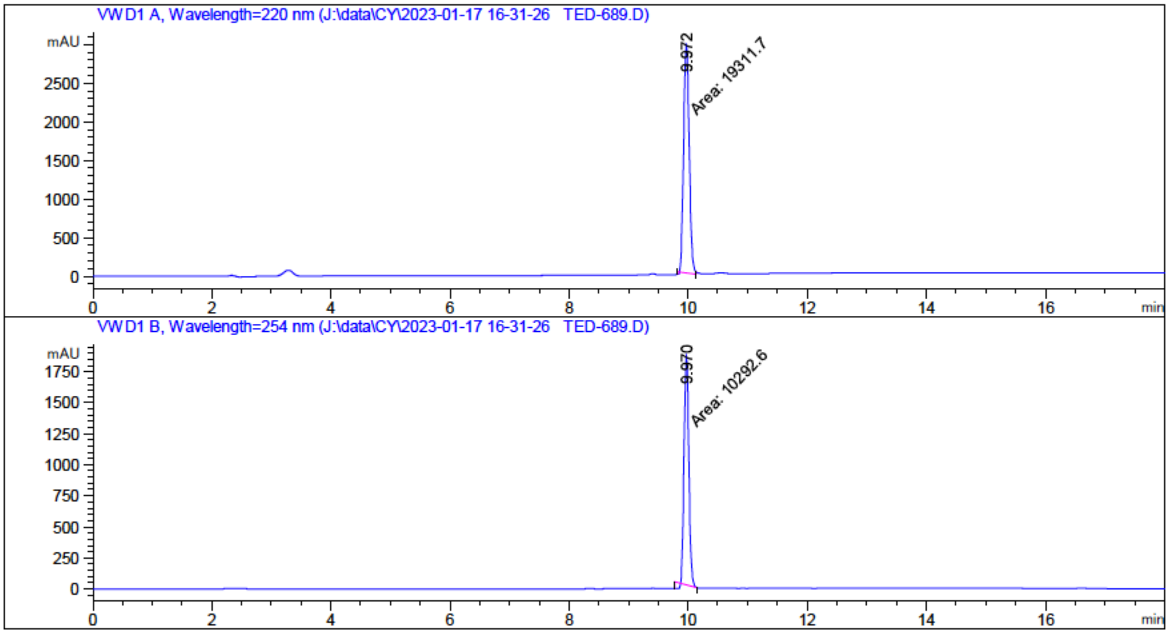


---

**UV 220 nm:**

**RT: 9.972 min Area %: 100.0000 %**

**UV 254 nm:**

**RT: 9.970 min Area %: 100.0000 %**

---

#### HPLC TED-690

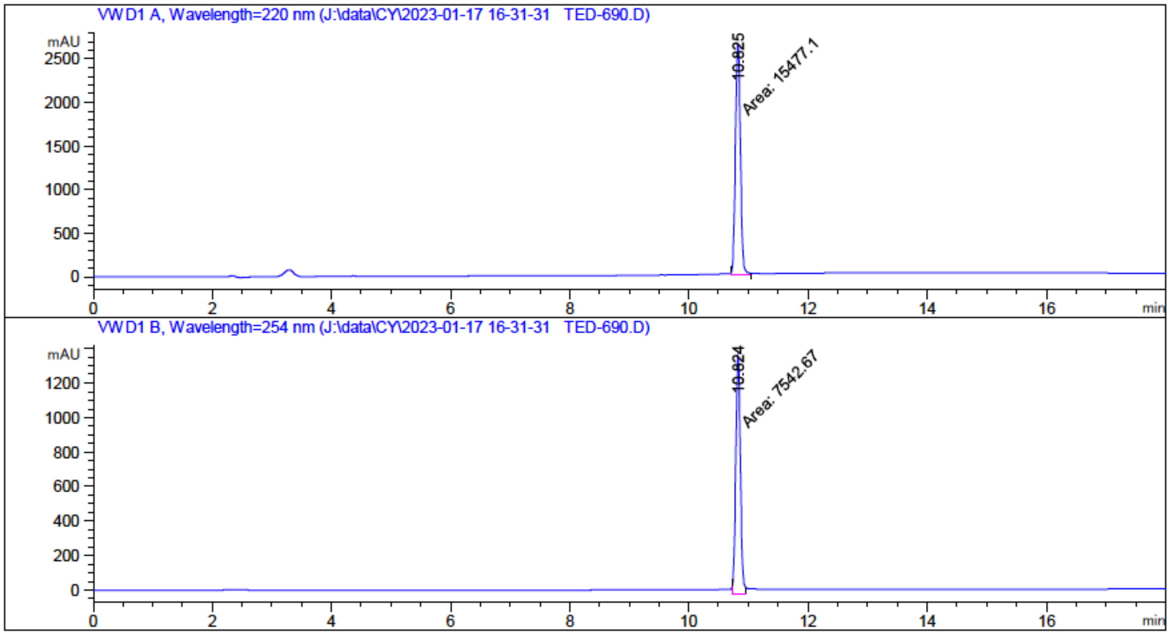


---

**UV 220 nm:**

**RT: 10.825 min** **Area %: 100.0000 %**

**UV 254 nm:**

**RT: 10.824 min** **Area %: 100.0000 %**

---

#### HPLC TED-734

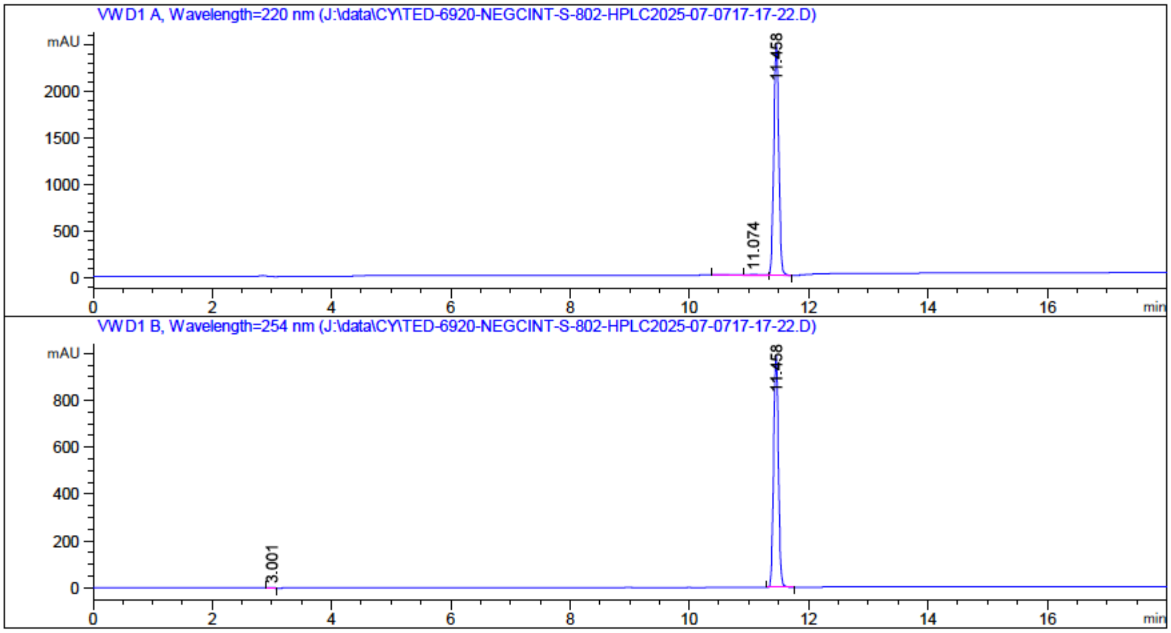


---

**UV 220 nm:**

**RT: 11.074 min** **Area %: 0.9026 %**

**RT: 11.458 min** **Area %: 82.1328 %**

**RT: 18.771 min** **Area %: 16.9646 %**

**UV 254 nm:**

**RT: 3.001 min** **Area %: 0.1760 %**

**RT: 11.458 min** **Area %: 96.5651 %**

**RT: 18.769 min** **Area %: 3.2589 %**

---
